# A CRISPR Screen Identifies Myc-associated Zinc Finger Protein (MAZ) as an Insulator Functioning at CTCF boundaries in *Hox* Clusters

**DOI:** 10.1101/2020.08.25.267237

**Authors:** Havva Ortabozkoyun-Kara, Pin-Yao Huang, Hyunwoo Cho, Varun Narendra, Gary Leroy, Jane A. Skok, Aristotelis Tsirigos, Esteban O. Mazzoni, Danny Reinberg

**Affiliations:** Howard Hughes Medical Institute, New York University School of Medicine, New York, New York 10016, USA; Department of Biochemistry and Molecular Pharmacology, New York University School of Medicine, New York, New York 10016, USA; Department of Pathology, New York University School of Medicine, New York, NY 10016, USA; Applied Bioinformatics Laboratories, New York University School of Medicine, New York, NY, USA; Department of Radiation Oncology, New York University School of Medicine, New York, NY, USA; Department of Medicine, Memorial Sloan Kettering Cancer Center, New York, NY, USA; Institute for Computational Medicine, NYU School of Medicine, New York, NY 10016; Department of Biology, New York University, New York, NY 10003, USA

## Abstract

The essential CCCTC-binding factor (CTCF) is critical to three-dimensional (3D) genome organization. CTCF binding insulates active and repressed genes within the *Hox* clusters upon differentiation, but such binding does not participate in boundary formation in all cell types, such as embryonic stem cells. We conducted a genome-wide CRISPR knockout screen to identify genes required for CTCF boundary activity at the *HoxA* cluster, complemented by novel biochemical approaches. This screen identified Myc-associated zinc finger protein (MAZ) as a CTCF insulator co-factor, among other candidates listed herein. MAZ depletion or specific deletion of MAZ motifs within the *Hox* clusters led to de-repression of posterior *Hox* genes immediately after CTCF boundaries upon differentiation, which phenocopied deletion of the proximal CTCF motifs. Similar to CTCF, MAZ interacted with the cohesin subunit, RAD21. Upon loss of MAZ, local contacts within topologically associated domains (TADs) were disrupted, as evidenced by HiC analysis. Thus, MAZ is a novel factor sharing insulation properties with CTCF and contributing to the genomic architectural organization.

**One Sentence Summary:** MAZ is identified as an insulator functioning at CTCF boundaries delimiting active and repressed genes at *Hox* clusters

## Main Text

Beyond the DNA sequence, chromatin structure and spatial organization of the genome regulate transcriptional output. Genomes of higher eukaryotes must be tightly folded and packaged within the nucleus (*1*). The partitioning of the genome into independent chromatin domains occurs via insulators. While several insulators are present in *Drosophila* (*2*), CTCF is the main insulator protein in vertebrates (*3–5*). CTCF is a highly conserved, ubiquitously expressed, eleven-zinc finger protein (*6*) that is critical for development (*7, 8*), and enriched at the borders of topologically associating domains (TADs) (*9–11*). Amongst the many proteins associated with CTCF at different loci (*4, 12*), only cohesin co-localizes to most CTCF binding sites and is required for CTCF function (*13, 14*). CTCF boundary activity is context-dependent (*15*). CTCF functions to form a strong boundary between active and repressed chromatin domains, decorated by Trithorax and Polycomb, at *Hox* clusters upon differentiation of mouse embryonic stem cells (mESCs) into cervical motor neurons (MNs). In general, CTCF-mediated boundaries thwart RNA polymerase II progression from active to repressed genes, thereby partitioning these antagonistic chromatin domains. Although mESCs are devoid of such chromatin barriers at *Hox* clusters, *Hox* gene boundaries clearly exhibit CTCF and cohesin occupancy (*16, 17*). Thus, during differentiation, additional regulatory factors appear necessary to foster CTCF-mediated insulation properties. We devised an unbiased genome-wide loss-of-function genetic screen involving a functional CTCF boundary within the *HoxA* cluster in cervical MNs.

We constructed a dual reporter ESC line (*Hoxa5:a7* mESCs) containing fluorescent reporters of *Hox* gene expression on each side of the CTCF demarcated boundary in the *HoxA* cluster by using CRISPR technology (*18*) (**Fig. 1A**, and **Fig. S1A**). The relative expression of *Hox* genes can be assayed in single cells, and any activity mediating CTCF-boundary formation can be assessed in this unique *Hoxa5:a7* dual-reporter system. As expected based on previous studies (*16, 19, 20*), *Hoxa5-P2A-mCherry* reporter expression was induced during cervical MN differentiation, while *Hoxa7-P2A-eGFP* remained repressed (**Fig. S1B-D**). To confirm that *Hoxa7-P2A-eGFP* reported defects in CTCF-dependent boundary formation, we deleted CTCF binding sites between *Hoxa5* and *Hoxa7* genes in ESCs (CTCF (Δ5|6) or CTCF (Δ5|6:6|7)) and demonstrated the de-repression of *Hoxa7-P2A-eGFP* by FACS analysis and RT-qPCR, as previously reported (**Fig. S1B-D**) (*16*). The ~10-15% *Hoxa7-P2A-eGFP* positive cells (**Fig. S1**), allowed for enough dynamic range to identify mutants that decrease or increase CTCF insulating properties.

**Fig. 1.**
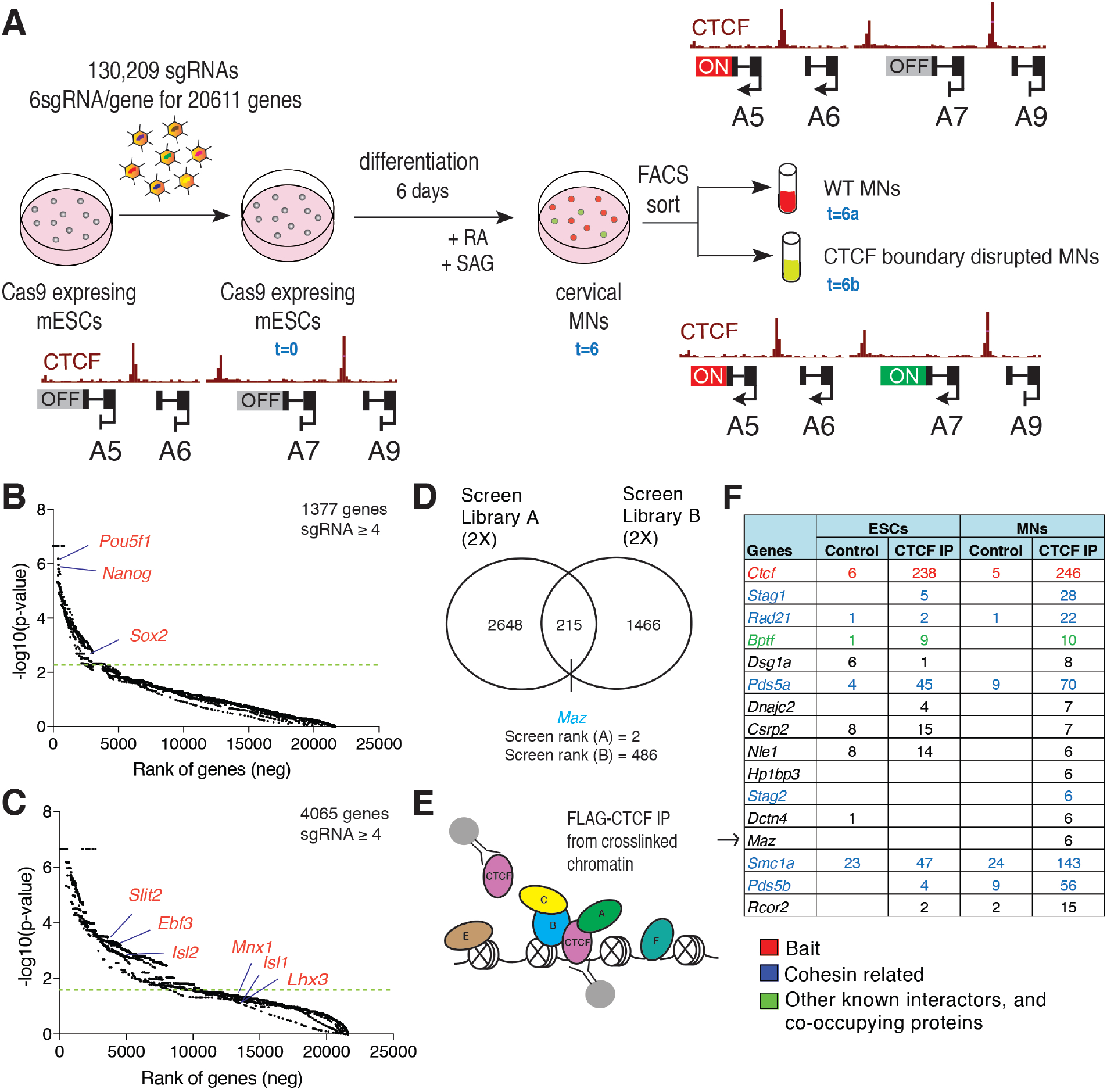
Genome-wide CRISPR loss-of-function screen to identify factors that affect the insulator function of CTCF, complemented with biochemical approaches. (**A**) Layout of the genetic loss-of-function screen that separates CTCF-boundary disrupted MNs from those with an intact boundary. (**B**) Rank of genes underrepresented in ESCs compared to plasmid library. Cut-off line indicates FDR < 0.05. (**C**) Rank of genes underrepresented in MNs compared to ESCs. Cut-off line indicates FDR < 0.05. (**D**) Venn diagram showing the overlap of CTCF-boundary related candidates identified in 4 independent (2 Library A, and 2 Library B) screens. P-value cut-off=0.05. (**E**) Scheme of crosslinked FLAG-CTCF ChIP-MS indicating identification of adjacent proteins on chromatin in addition to interactors. (**F**) Crosslinked FLAG-CTCF ChIP-MS in ESCs and MNs results in identification of known CTCF interactors, and novel proteins. The peptide counts in FLAG-CTCF IPs were normalized to control FLAG IPs in untagged cells. The list is ranked based on CTCF IP/control ratios in MNs.

To identify factors required for the integrity of the CTCF-mediated boundary, we performed an unbiased loss-of-function genetic screen on *Hoxa5:a7* dual reporter ESCs using a pooled genome-wide library of single-guide RNAs (sgRNAs) (*21*), as demonstrated in **Fig. 1A**. A *Hoxa5:a7* ESC clone expressing *Cas9* (**Fig. S2A**) was transduced with the pooled lentiviral sgRNAs at a low multiplicity of infection (MOI) of ~0.4 as applied before (*21, 22*), such that each transduced cell expressed a single sgRNA. The reporter ESCs (t=0) were then differentiated into cervical MNs (t=6) via the addition of RA/SAG (*23*), and sorted by FACS into two MN populations: **(1)** WT MNs (*mCherry* positive/*eGFP* negative cells), t=6a and **(2)** CTCF boundary disrupted MNs (double positive cells), t=6b. By preparing libraries at each time-point, the relative sgRNA representation at t=0, 6, 6a, and 6b were compared using next generation sequencing, as described previously (*21, 22, 24, 25*). This screen setup enabled identification of three sets of genes: (1) Essential genes in mESCs (negative selection), (2) Essential/differentiation related genes in MNs (negative selection), and (3) Genes affecting CTCF-boundary function (positive selection). Throughout the screen, *Hox* gene reporter expression was not observed in ESCs upon perturbation of any gene, which increased our confidence in identifying CTCF-boundary related candidates rather than general repressors.

As expected from a functional screen, we observed selective loss of essential genes in the starting population (ESCs, t=0) compared to plasmid library (**Fig. 1B**, and **Fig. S2B-C**), and further loss of genes essential/required for MN differentiation (MN, t=6) compared to ESCs (t=0) population (**Fig. 1C**, and **Fig. S2D-E**), indicating the success of the screen. Among genes underrepresented in MNs compared to ESCs (FDR<0.05), we observed Polycomb group genes, CTCF, cohesin components, and several components related to the MN differentiation pathway. Our genome-wide screens were performed in duplicates by using independent genome-wide sub-libraries (library A, and library B), resulting in four independent screens. In each screen, we identified ~1000 genes positively selected in CTCF-boundary disrupted MNs (t=6b) compared to WT MNs (t=6a) by using the Model Based Analysis of Genome-wide CRISPR Knock-out Screen (MaGeCK) tools (*26, 27*). We narrowed the candidate list to 215 genes in CTCF-boundary disrupted MNs compared to WT MNs out of four independent sub-library screens, **Fig. 1D**. Notably, *Maz* was identified as a top candidate (rank=2) in one of the genome-wide screens, and also detected in similar screen (rank=486) (**Fig. 1D** and **Data S1**).

We complemented the genetic screen with orthogonal biochemical approaches for the identification of proteins co-localizing with CTCF on chromatin. Unlike previous studies to identify CTCF partner proteins in soluble cellular fractions through the use of overexpression based systems (*12, 28*), we identified proteins co-localizing with CTCF on chromatin that may or may not interact with CTCF, but nonetheless may be important for its insulation properties *in situ*. To pull-down CTCF under endogenous conditions, we generated an ESC line containing C-terminal FLAG-tagged CTCF via CRISPR technology (**Fig. S3A**) (*18*), and confirmed successful FLAG-CTCF immunoprecipitation (IP) from the nuclear fraction of ESCs (**Fig. S3B**, and see **Fig. S3D**). To expand and identify factors co-localizing with CTCF on chromatin, we applied two biochemical methods: (1) FLAG-CTCF immunoprecipitation (IP) from native chromatin in ESCs and MNs (**Fig. S3C**), (2) FLAG-CTCF immunoprecipitation (IP) from crosslinked chromatin in ESCs and MNs (**Fig. 1E**), an adapted version of the chromatin immunoprecipitation (ChIP)-mass spectrometry (ChIP-MS) approach described previously (*29–32*). Affinity purification performed here enriched for CTCF-bound chromatin fragments using FLAG pull-down followed by FLAG peptide elution to minimize nonspecific interactions. We then performed mass spectrometry (MS) on the eluted proteins. In both ESCs and MNs, wild-type cells without FLAG-tag acted as background controls to normalize FLAG IPs. In both Flag-CTCF ChIP-MS approaches, we identified known interactors and novel proteins interacting or co-binding with CTCF (**Fig. 1E-F**, **Fig. S3C**, and see **Data S2** for all candidates). As expected, we recovered cohesin components and accessory subunits, and other chromatin remodelers (**Fig. 1F** and **Fig. S3C**). Interestingly, MAZ was identified uniquely in the crosslinked based CTCF ChIP-MS and served as an overlapping candidate based on the *Hox* related functional screen and CTCF ChIP-MS approach. Thus, Maz might serve a role in regulating the CTCF-boundary at the *Hox* loci. MAZ was also reported as co-localizing with CTCF at ~48% of binding sites based on ENCODE ChIP sequencing (ChIP-seq) data in K562 cells (*33*), reinforcing our observation of their proximal binding on the cross-linked chromatin.

Both genetic and biochemical approaches revealed a large list of candidates, which were further narrowed down and validated through independent secondary genetic screens. In order to systematically narrow down candidates from the primary genome-wide screens (**Data S1**), and check whether CTCF partners identified in **Fig. 1F** and **Fig. S3C** (**Data S2**) have a role at the CTCF-boundary at the *HoxA* cluster, we performed secondary loss-of-function screens with a small custom library (**Data S3**). This custom library included sgRNAs targeting the list of candidates from primary screens, and other proteins that co-purified with CTCF (**Fig. S4E**). The sgRNAs in the custom library were retrieved from another genome-wide library constructed with improved design tools (*34*). Importantly, these secondary screens were performed with increased statistical power in ESCs having either the wild-type *Hoxa5:7* reporter (**Fig. 2A**, and **Fig. S4A-B**), or the CTCF (Δ5|6:6|7) *Hoxa5:7* reporter (**Fig. 2D**, and **Fig. S4C-D**) to focus on candidates uniquely impacting the CTCF-boundary in the wild-type background. Based on the rank of genes overrepresented in the *Hoxa5:7* dual positive MN population compared to *Hoxa5* positive cells, we identified 55 genes that disrupt the CTCF-boundary in the wild-type background having intact CTCF binding sites (**Fig. 2B**, **Data S4**). Similarly, we identified 165 genes that influence the CTCF-boundary/*Hoxa7* gene expression from screens performed in the CTCF (Δ5|6:6|7) background (**Fig. 2E**, **Data S5**). Thus, the secondary screens resulted in a small gene list (43 genes) that when mutated phenocopied CTCF (Δ5|6) motif deletion in the presence of intact CTCF binding sites (see **Fig. S4F** for comparison of secondary screens in both backgrounds, **Data S6**). Importantly, the secondary screens also confirmed the identification of *Maz* uniquely in the wild-type background. Other genes shown in **Fig. 2B** and **Fig. 2E** are expected positive controls such as *Ctcf*, cohesin components/accessory subunits, and *Znf143* that is reported to co-localize with CTCF at TADs (*35*).

**Fig. 2.**
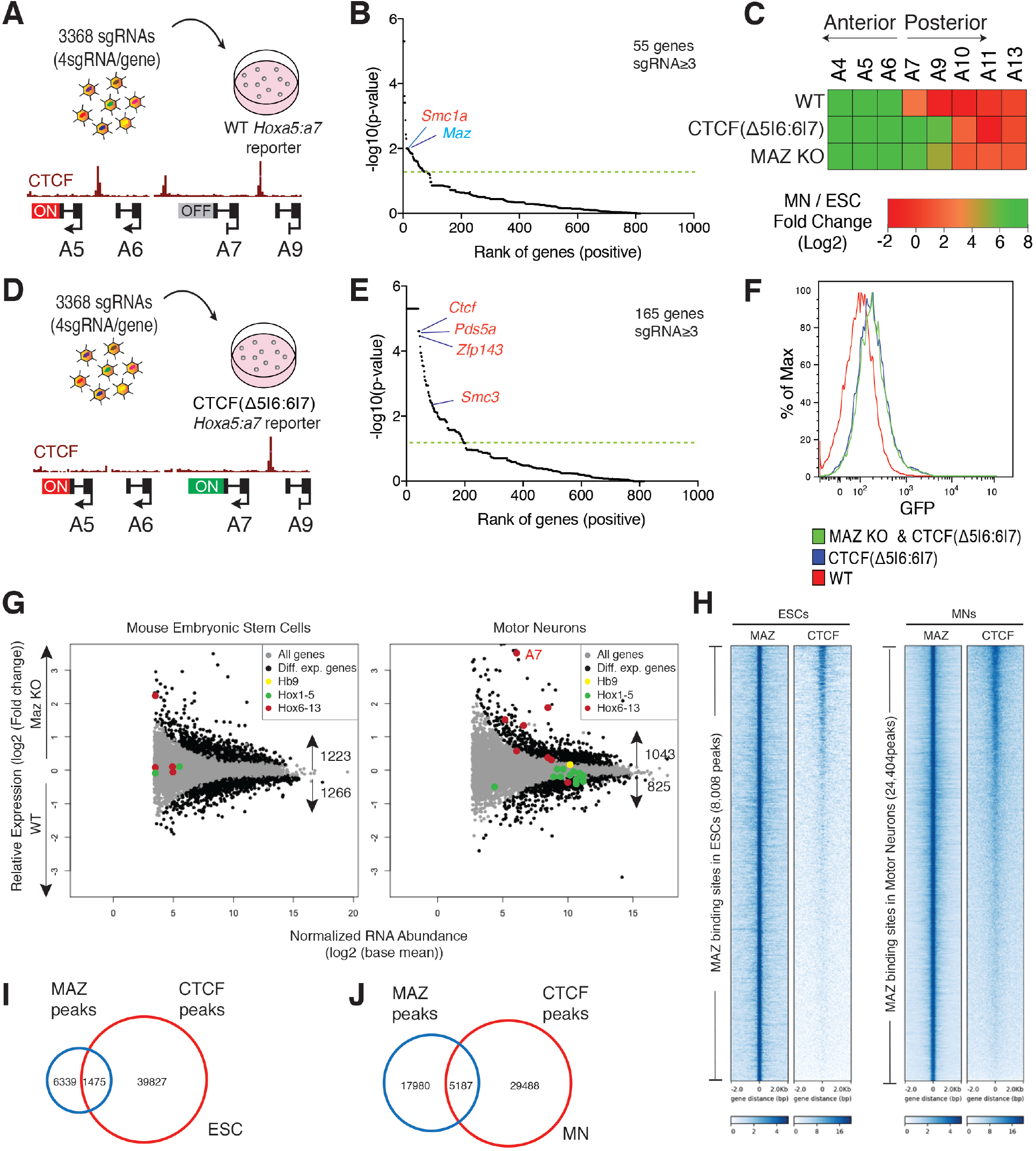
Secondary CRISPR loss-of-function screens, and individual validation of MAZ as an insulator functioning at CTCF-boundaries in *Hox* clusters. (**A**) Scheme of secondary screen performed in WT background. (**B**) Rank of genes overrepresented in boundary disrupted MNs versus WT MNs in one biological replicate. Cut-off line indicates p < 0.05. (**C**) Heat map of relative gene expression in WT, CTCF(Δ5|6:6|7), and MAZ KO at the *HoxA* cluster in MNs versus ESCs from three biological replicates. Maz KO represents three independent clones. (**D**) Scheme of secondary screen performed in CTCF (Δ5|6:6|7) background. (**E**) Rank of genes overrepresented in dual positive *Hoxa5:a7* MN population (further disrupted boundary) versus *Hoxa5-mCherry* positive population (WT) in two biological replicates. Cut-off line indicates p < 0.05. (**F**) Flow cytometry analysis of MNs with the indicated genotypes: WT, CTCF(Δ5|6:6|7), and MAZ KO & CTCF(Δ5|6:6|7). This plot is one representation of three biological replicates quantified in **Fig. S6G**. (**G**) RNA-seq MA plot of WT versus MAZ KO ESCs (left), and MNs (right) from three biological replicates. Differentially expressed genes are selected as p-value adjusted (padj) < 0.05. *Hox* genes in 4 *Hox* clusters are colored based on their position with respect to the previously demonstrated CTCF-boundary in MNs. *Hb9* is a MN marker. (**H**) Heat maps of CTCF and MAZ ChIP-seq read density in ESCs and MNs within a 4 kb window centered on the maximum value of the peak signal. (**I** to **J**) Overlap of CTCF and MAZ binding sites in ESCs and MNs. ChIP-seq experiments are from one representative of two biological replicates.

Among the candidates identified to mimic CTCF (Δ5|6) at the *HoxA* cluster, MAZ has been ranked high in multiple primary screens, identified as a co-localizing factor with CTCF on chromatin, and further validated through secondary screens. MAZ is a ubiquitously expressed protein, initially identified as a regulatory protein associated with *Myc* gene expression (*36*), and also identified as a regulatory factor for the insulin promoter (*37*). To validate the screen results, we generated MAZ knock-out (KO) in mESCs through CRISPR (**Fig. S5A-B**). MAZ KO did not produce a profound change in gene markers associated with ESC and MN fate (**Fig. S6D**). In addition, MAZ KO did not result in cell cycle changes in ESCs (**Fig. S6E-F**). MAZ KO did not affect overall CTCF and cohesin levels (**Fig. S5B**). As shown in **Fig. 2C**, MAZ KO in MNs mimicked the specific deletion of the CTCF site (Δ5|6:6|7) at the *HoxA* cluster and disrupted the boundary between active and repressed genes. In addition, MAZ KO resulted in differential expression of ~2400 genes in ESCs (**Fig. 2G**, left; see **Fig. S5C** for GO analysis and **Data S7**), and ~1800 genes in MNs compared to WT (**Fig. 2G**, right and **Data S8**). Gene Ontology (GO) analysis indicated that developmental processes, particularly anterior-posterior pattern specification, are enriched in MAZ KO MNs compared to WT MNs (**Fig. S5D**). Consistent with a boundary role and with CTCF binding site deletions, MAZ KO led to de-repression of posterior *Hox* genes in MNs, but not in ESCs (**Fig. 2G**, see RT-qPCR data in **Fig. S6A-C**). We did not observe further de-repression of *Hoxa7-GFP* upon differentiation of CTCF (Δ5|6:6|7) mESCs having a *Maz* deletion into MNs (**Fig. 2F**, see **Fig. S6G** for quantification), confirming our secondary screen results (**Fig. 2D-E**).

Based on our ChIP-seq analysis, ~20% of MAZ binding sites co-localize with CTCF in mESCs and MNs (**Fig. 2H-J)**. MAZ mostly binds to promoters in addition to introns, intergenic regions, and other regions (**Fig. S7A**). As we initially identified the MAZ KO as influencing the CTCF-boundary at the *HoxA* cluster (**Fig. 2C**), we compared ChIP-seq tracks of MAZ at the *HoxA* cluster and to those of CTCF. We observed co-localization of MAZ and CTCF at CTCF borders in *Hox* clusters as shown for the *HoxA* cluster in **Fig. 3A** (**Fig. S9A-B** for *HoxD*, and **Fig. S10A-B** for *HoxC*). As shown in **Fig. 3A**, MAZ KO in ESCs and MNs resulted in a slight decrease of CTCF binding at the boundary at *HoxA* cluster. We also observed a similar global decrease in CTCF binding in MAZ KO (**Fig. S7B-C**). Although MAZ and CTCF co-localize on the cross-linked chromatin (**Fig. 1F**, and **Fig. 2H**), they did not co-immunoprecipitate (co-IP) under native conditions (**Fig. S3D-F**). Interestingly, we observed that similar to CTCF, MAZ co-IPs with RAD21, the cohesin component (**Fig. 4D**, and **Fig. S3E-F**). Therefore, MAZ appears to interact with the cohesin component, and bind to DNA in the CTCF proximity.

**Fig. 3.**
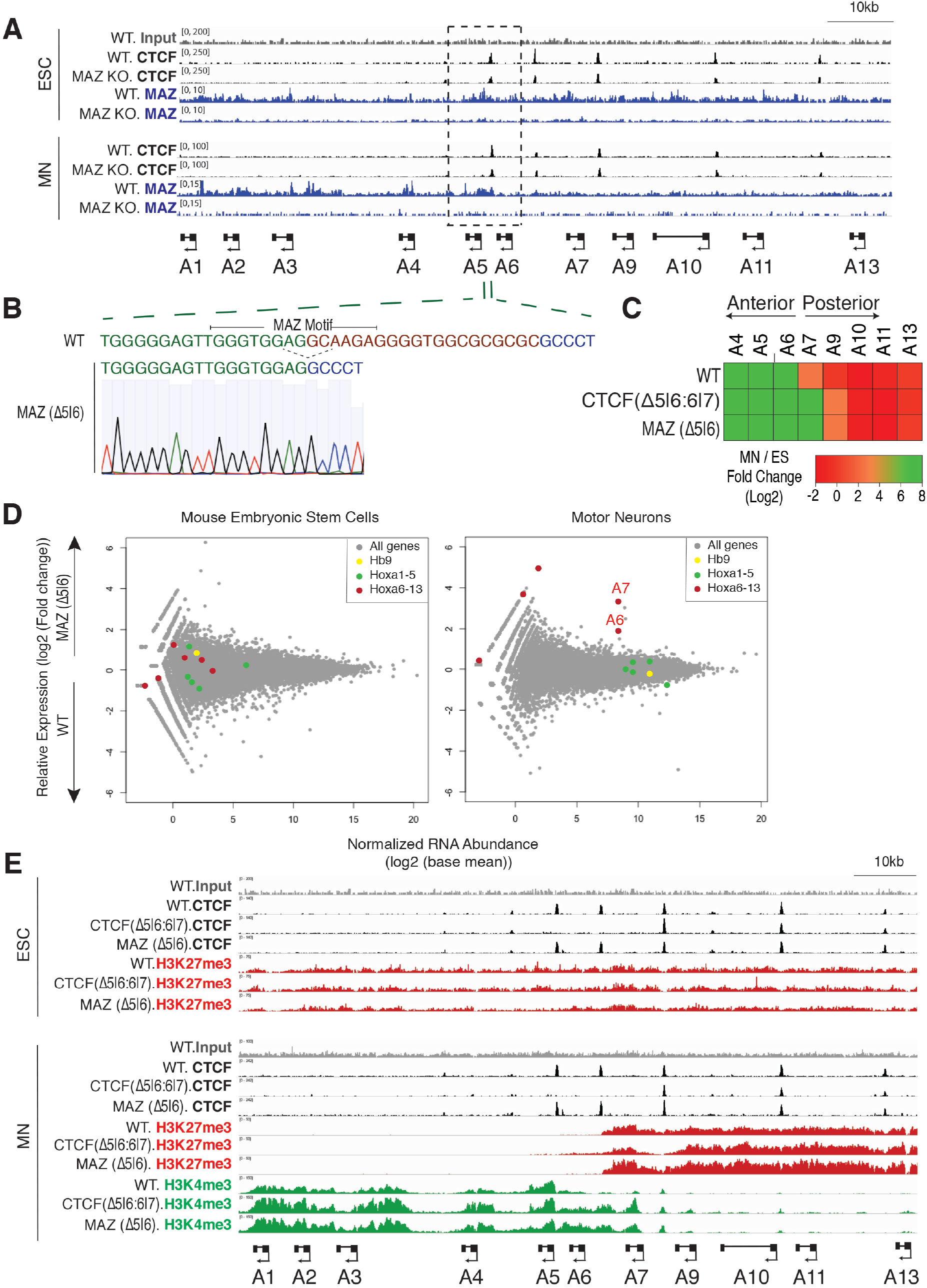
Loss of MAZ binding site alters *Hox* gene expression pattern and chromatin domains at *Hox* clusters. (**A**) ChIP-seq for CTCF or MAZ in WT and MAZ KO ESCs and MNs in the *HoxA* cluster. (**B**) MAZ binding site deletion via CRISPR is depicted for the 5|6 site at the *HoxA* cluster. (**C**) Heat map of relative gene expression in WT, CTCF(Δ5|6:6|7), and MAZ (Δ5|6) at the *HoxA* cluster in MNs versus ESCs from three biological replicates. (**D**) RNA-seq MA plot of WT versus MAZ (Δ5|6) ESCs (left), and MNs (right) from three biological replicates. *HoxA* genes are colored based on their position with respect to the previously demonstrated CTCF boundary in MNs. *Hb9* is a MN marker. (**E**) ChIP-seq for CTCF, and indicated histone modifications in WT, CTCF (Δ5|6:6|7), and MAZ (Δ5|6) ESCs and MNs in the *HoxA* cluster. ChIP-seq tracks are from one representative of two biological replicates for CTCF, one replicate for the histone modifications.

**Fig. 4.**
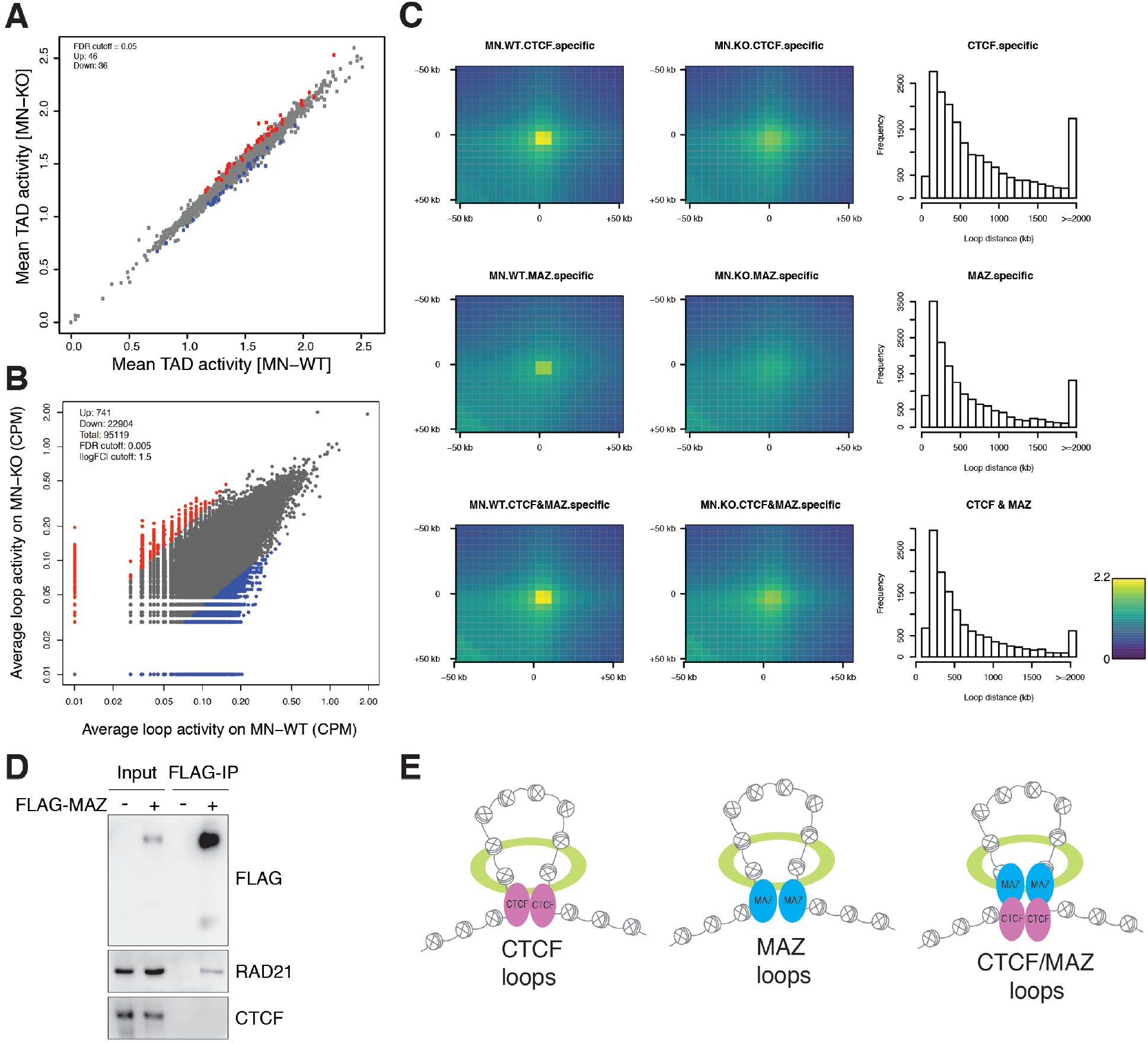
Effect of MAZ on global genome organization. (**A**) Scatter plot showing differential intra-TAD activity in WT vs MAZ KO MNs (FDR cut-off=0.05). (**B**) Scatter plot showing differential loop activity in WT vs MAZ KO MNs (all loops, n=95119, FDR cut-off=0.005, | log (Fold Change) | cut-off=1.5, Up-regulated=741, Down-regulated: 22904). (**C**) Aggregate Peak Analysis (APA) of loops in WT vs MAZ KO MNs showing ChIP-seq signals of CTCF, MAZ, or both at any region covered by them. The resolution of APA is 5 kb. Histograms showing the distribution of loop distance in MAZ KO compared to WT related to the binding level of ChIP-seq. (**D**) Western blot analysis of FLAG, RAD21, and CTCF upon FLAG-MAZ immunoprecipitation from nuclear extract of 293FT cells. (**E**) Proposed model for the effect of MAZ on loops via its interaction with cohesin (middle) or binding adjacent to CTCF and interaction with cohesin (right), in addition to existing loop-extrusion model of CTCF and cohesin (left, (*44, 45*)).

MAZ binds to a GC-rich motif on DNA (GGGAGGG) through its zinc fingers (**Fig. S7D-E**) (*38*). We found MAZ binding motifs close to CTCF at *Hox* boundaries and tested whether deletion of MAZ binding motifs at a specific *Hox* cluster mimics deletion of a CTCF binding site in the respective *Hox* cluster. We reasoned that if MAZ binding close to a CTCF boundary has a direct effect on anterior-posterior patterning, deletion of its binding motif could alter *Hox* gene expression similar to MAZ KO in MNs. Interestingly, MAZ binding site deletions generated at *Hoxa5*|*6* (**Fig. 3B**), *Hoxd4*|*8* (**Fig. S9**), and *Hoxc5*|*6* (**Fig. S10**) phenocopied CTCF binding site deletions at the respective boundaries without altering CTCF binding at the boundary (**Fig. 3C-E**, **Fig. S8C**, and **Data S9-10** for *HoxA;* **Fig. S9A** and **Fig. S9E** for *HoxD;* and **Fig. S10A** and **Fig. S10E** for *HoxC* clusters). As expected, MAZ binding site deletions at *Hox* clusters did not influence the cell cycle in ESCs (**Fig. S8A-B** for *HoxA*, **Fig. S9C-D** for *HoxD*, and **Fig. S10C-D** for *HoxC* clusters). These results pointed to a specific role of MAZ binding in regulating *Hox* gene expression at CTCF boundaries in multiple *Hox* clusters during differentiation. When we analyzed how chromatin domains were influenced upon deletion of the MAZ binding site at the *Hoxa5*|*6* boundary, we observed spreading of active chromatin (H3K4me3) into the repressed region (H3K27me3) at the boundary, similar to the CTCF binding site deletion shown in **Fig. 3E**, and to that previously reported (*16*) (see **Fig. S9A** and **Fig. S10A** for *HoxD* and *HoxC* clusters). Similar to MAZ (Δ5|6) not altering the neighboring CTCF binding or RAD21 binding (data not shown), CTCF (Δ5|6:6|7) did not affect adjacent MAZ binding at the *HoxA* cluster (**Fig. S7F**).

Besides its boundary role at *Hox* clusters, CTCF plays a pleiotropic role in 3D genome structure. MAZ co-localizes with CTCF at ~20% of MAZ binding sites in mESCs and MNs, and MAZ KO reduces CTCF binding. Thus, we examined the effect of MAZ KO in global genomic organization by performing HiC in WT versus MAZ KO ESCs and MNs in biological duplicates (**Fig. 4**, and **Fig. S11** to **S14**). MAZ depletion resulted in alteration of local contacts within topologically associated domains (TADs) in ESCs (**Fig. S11C**) and MNs (**Fig. 4A**), indicating the contribution of MAZ to the integrity of TADs. As expected, the active (A) and inactive (B) compartments were not affected upon MAZ KO in ESCs and MNs (**Fig. S11A**). There were also changes in boundary scores upon MAZ KO in ESCs and MNs as shown by PCA (**Fig. S11B**). In addition, analysis of differential loop activity showed alterations upon MAZ KO in both cell types (**Fig. 4B**, and **Fig. S11D)**, indicating the global loss of loops in MAZ KO compared to WT. Consistent with this observation, aggregate peak analysis (APA) showed that contact frequencies were decreased in MAZ KO ESCs (**Fig. S11E**), and MNs (**Fig. 4C**) compared to WT, and that these changes occurred in short ranges within 2 Mb. As CTCF is known to be present at loop anchors, we hypothesized that there might be loops bound by MAZ alone or MAZ and CTCF together. Therefore, we categorized the loops based on CTCF, MAZ, or MAZ and CTCF occupancy, and investigated how they were affected upon MAZ KO. Interestingly, the loops specific to CTCF, MAZ, or MAZ and CTCF showed a decrease in contact frequencies upon MAZ KO (**Fig. 4C**, and **Fig. S11E**). Moreover, we observed that some loop domains in larger ranges are also slightly downregulated upon MAZ KO while we did not observe local effects of MAZ KO on *Hox* cluster organization with respect to the CTCF and MAZ boundary, within the resolution of HiC (**Fig. S12** to **S14**). While we did not detect interaction between MAZ and CTCF (**Fig. S3E-F**), MAZ and cohesin components did exhibit interaction (**Fig. 4D**), as is the case with CTCF (**Fig. S3E-F**, and (*12*)). Moreover, as deletion of either the MAZ or CTCF binding motif results in spreading of an active domain into a repressed one (**Fig. 3C-E**, and (*16*)), we speculate that MAZ, CTCF and cohesin function together to thwart progression of transcription from the respective active domain.

These results demonstrate that an unbiased genome-wide CRISPR screen coupled with biochemical approaches enable identification of novel factors that function similar to and in conjunction with CTCF. Our results place MAZ as a novel factor that functions in partitioning *Hox* clusters into insulated domains, wherein Trithorax and Polycomb activities are important to maintain distinct *Hox* gene expression. MAZ binding site deletions at active and repressed gene borders in *Hox* clusters phenocopy the effect of CTCF binding site deletions at *Hox* clusters (*16, 39*). Consistent with our findings, *Maz* ^−/−^ mice show perinatal lethality, and developmental defects in the kidney and urinary track (*40*), though other phenotypes remain to be tested. Although more than 25 out of 39 *Hox* genes are expressed in the kidney, the role of *Hox* genes other than the *Hox11* paralogous group seems to be under active investigation (*41*). Hence, it can be informative to study possible *Hox* gene misexpression patterns in the kidney or urinary track in *Maz* ^−/−^ mice. MAZ contributes to the integrity of TADs, and contacts within TADs and loops are impacted upon loss of MAZ although the effects are not as large-scale as the loss of essential architectural proteins such as CTCF (*42*) or cohesin (*43*). In particular, MAZ appears to be present at the loop anchors with or without CTCF, and both proteins interact with cohesin components independently (**Fig. 4E**). Based on our preliminary transcription assays, MAZ and CTCF binding impact the progression of transcription in the presence of cohesin. Taken together, our findings point to MAZ functioning as a novel insulator factor at *Hox* clusters and being critical to genome organization. In addition to its role in impacting loop interactions, MAZ, together with adjacent CTCF and cohesin, possibly functions by blocking RNA polymerase II progression.

## Acknowledgments

We thank L. Vales for reading and guidance on the manuscript. We thank S. Tu, S. Krishnan, and T. Escobar for discussions; O. Oksuz for discussions regarding sequencing data analysis; Y. Grobler for providing *Drosophila* S2R+ cells; and D. Hernandez for technical assistance. We thank other past and present members of the Reinberg laboratory for discussions as the work was in progress. We also thank New York University Langone Medical Center (NYULMC) Genome Technology Center, particularly A. Heguy, P. Zappile, and P. Meyn, for help with sequencing, the Applied Bioinformatics Laboratories (ABL) for providing bioinformatics support, and NYULMC Cytometry and Cell Sorting Core, particularly P. Lopez, and M. Gregory for help with FACS. This study utilized computing resources at the High-Performance Computing Facility of the Center for Health Informatics and Bioinformatics at the NYULMC.

## Author contributions

H.O.K., E.O.M., and D.R. conceived the project, designed the experiments, and wrote the paper; H.O.K. performed most of the experiments and the bioinformatic analysis except HiC; P.Y.H. helped with immunoprecipitation, transcription constructs, and RT-qPCRs; H.C. performed the bioinformatic analysis of HiC under the supervision of A.T.; V.N. advised in the initial design of study; G.L. advised on the mass-spectrometry and performed preliminary transcription assays; and J.A.S. advised on the progression of this study.

## Data and materials availability

Sequencing data has been deposited at Gene Expression Omnibus and accession numbers will be available immediately upon publication.

## Supplementary Materials

### Materials and Methods

#### Cell Culture and Motor Neuron Differentiation

E14 mouse embryonic stem cells (mESCs) were cultured in standard medium supplemented with LIF, and 2i conditions (1 mM MEK1/2 inhibitor (PD0325901, Stemgent) and 3 mM GSK3 inhibitor (CHIR99021, Stemgent)). For motor neuron (MN) differentiation, the previously described protocol was applied (*16*). Briefly, ESCs were differentiated into embryoid bodies in 2 days, and further patterning was induced with addition of 1 μM all-trans-retinoic acid (RA, Sigma) and 0.5 μM smoothened agonist (SAG, Calbiochem). Biological replicates stand for independent differentiation experiments performed. 293FT cells were cultured in standard medium as described in the manufacturer’s protocol (Thermo Fisher Scientific).

#### CRISPR Genome Editing

sgRNAs were designed using CRISPR design tools in http://crispr.mit.edu/, currently available in https://benchling.com. All sgRNAs were cloned into SpCas9-2AGFP vector (Addgene: PX458) or into a lentiviral vector lentiGuide-puro (Addgene: 52963). The sgRNAs were transfected into mESCs using Lipofectamine 2000 (Invitrogen), as described before (*16*) or infected into a lentiCas9-blast (Addgene: 52962) expressing mESC clone. In the case of CRISPR knock-in cell lines, donor DNA (1 μl of 10 μM single stranded DNA oligo, or 3 μg pBluescriptSK (+) plasmid containing donor DNA) were transfected with 1 μg px458-sgRNAs. Single clones from GFP-positive FACS sorted cells or puromycin (InvivoGen)-resistant cells were genotyped and confirmed by sequencing. When necessary, PCR products were further assessed by TOPO cloning, and sequencing to distinguish the amplified products of different alleles. The sequencing chromatograms were aligned in Benchling. All sgRNAs, donors, and genotyping primers are shown in **Table S1**.

#### Cell Line Generation

##### *Hoxa5:a7* dual reporter in WT and CTCF (Δ5|6) backgrounds

To generate *Hoxa5:a7* dual reporter cells, mESCs were sequentially targeted at *Hoxa5* and *Hoxa7* loci, respectively. mESCs were initially transfected with sgRNA and donor pBluescriptSK (+) plasmid for *Hoxa5-P2A-mCherry* cell line generation using Lipofectamine (Invitrogen). *Hoxa5-mCherry* cell line was confirmed through genotyping, sequencing, and FACS analysis upon MN differentiation for the homozygous insertion of reporter. Next, the *Hoxa5-mCherry* cell line was transfected with sgRNA and donor pBluescriptSK (+) plasmid for generation of the dual *Hoxa5:a7* knock-in cell line, which was confirmed by genotyping, sequencing, and FACS analysis for the homozygous insertion of reporter. To demonstrate *Hoxa7-P2A-GFP* expression in MNs, CTCF binding sites at *Hoxa5*|*6* and *Hoxa6*|*7* were removed via sequential CRISPR genome editing using respective sgRNAs, generating CTCF (Δ5|6) and CTCF (Δ5|6:6|7) deletion lines in the *Hoxa5:a7* dual reporter background. For CRISPR library screen experiments, WT or CTCF (Δ5|6:6|7) dual reporter lines were transduced with lentiCas9-blast (Addgene, 52962), and *Cas9* expressing clones were obtained after selection with blasticidin (InvivoGen).

#### FLAG-CTCF tagged cell line

To generate the CTCF C-terminal FLAG-tagged cell line, E14 ESCs were targeted with sgRNA in SpCas9-2AGFP vector (Addgene: PX458) and single-stranded donor oligo at the CTCF locus and the cell line was confirmed by genotyping, sequencing, and western blot for FLAG-CTCF.

#### MAZ KO cells

WT or CTCF (Δ5|6:6|7) *Hoxa5:a7* dual reporter cells expressing *Cas9* were targeted with sgRNAs in lentiGuide-puro vector for *Maz*. Knock-out of *Maz* was confirmed by genotyping, sequencing, and western blot.

#### MAZ binding site deletions

*Hoxa5:a7* dual reporter cells were targeted with sgRNAs in SpCas9-2AGFP vector (Addgene: PX458) for MAZ binding sites at *HoxA, HoxD* or *HoxC* cluster. Specific MAZ binding site deletions were confirmed by genotyping and sequencing.

#### CRISPR Screens

CRISPR genome-wide screens were done using methods described previously (*21, 22*). Briefly, GeCKO genome-wide pooled CRISPR libraries (Addgene: 1000000053) were amplified, and deep-sequenced to confirm sgRNA representations, as shown previously (*21*). A *Cas9* expressing *Hoxa5:a7* ESC clone was transduced with the pooled lentiviral sgRNAs at a low multiplicity of infection (MOI) of ~0.4. The reporter ESCs were selected with puromycin, cultured for 7 days, and differentiated into MNs in 6 days, and sorted by FACS into two MN populations: (1) WT MNs (*mCherry* positive/*eGFP* negative cells) and (2) CTCF-boundary disrupted MNs (double positive cells). During the screens, 300X and 1000X coverage was applied for genome-wide screens, and secondary screens, respectively. CRISPR libraries were prepared at each-time point and/or sorted population, and the relative sgRNA representation was assessed using next generation sequencing, as described previously (*21, 22*).

#### Custom Library Construction for Secondary CRISPR Screens

sgRNAs for custom library used in the secondary CRISPR screens were retrieved from a previously designed genome-wide mouse CRISPR knock-out pooled library (Brie) (*34*). When required for several genes, sgRNAs were designed by using the Broad Institute CRISPRko gRNA design tools (currently at https://portals.broadinstitute.org/gpp/public/analysis-tools/sgrna-design). All sgRNAs in the custom library in **Data S3** were synthesized as a pool by Twist Bioscience. The custom library was cloned into lentiGuide-puro vector, amplified, and verified in terms of representation of all constructs using the methods described previously (*46*).

#### Flow Cytometry

Cells were trypsinized, filtered, and stained with 4,6-diamidino-2-phenylindole (DAPI, Sigma) to eliminate dead cells during analysis of *Hoxa5:a7* reporters in ESCs and MNs. *Hoxa5:a7* dual fluorescent reporter cells in WT versus other backgrounds were assessed by using single color fluorescent reporters as controls in the same cell type as analyzed (*i.e.* MNs). *Hb9-T2A-GFP* reporter cells (not shown) were used as GFP control in MNs. For cell cycle analysis, ESCs were fixed in 75% Ethanol, and DNA was stained with Propidium iodide (Thermo Fisher Scientific) after RNase A (Thermo Fisher Scientific) treatment. FlowJo 8.7 was used for all FACS analysis.

#### Expression Analysis

RNA was purified from cells with RNAeasy Plus Mini kit (Qiagen), and reverse transcription was performed on 1 μg RNA by using Superscript III (Life Technologies) and random hexamers (Thermo Fisher Scientific). RT-qPCRs were performed in replicates on 100 ng cDNA using PowerUp SYBR Green Master Mix (Thermo Fisher Scientific). The primers are listed in **Table S2**. For RNAseq analysis, 1 μg RNA was used to prepare ribominus RNAseq libraries according to the manufacturer’s protocols by the NYU Genome Technology Center.

#### Chromatin Immunoprecipitation Sequencing

ChIP-seq experiments were performed as described (*47*) [see details about ESC fixation (*47*), and MN fixation in (*16*)]. Briefly, cells were fixed with 1% formaldehyde, nuclei were isolated, and chromatin was fragmented to ~250 bp in size using a Diagenode Bioruptor. ChIP was performed by using antibodies listed in **Table S2**. Chromatin from *Drosophila* (1:100 ratio to ESC or MN derived chromatin), and *Drosophila*-specific H2Av antibody were used as spike-in control in each sample. For ChIP-seq, libraries were prepared as described (*16*) using 1-30 ng of immunoprecipitated DNA. ChIP-qPCRs were performed with PowerUp SYBR Green Master Mix (Thermo Fisher Scientific) and detected by the Stratagene Mx3005p or QuantStudio 5 (Thermo Fisher Scientific) instrument. All ChIP-qPCR primers are listed in **Table S2**.

#### Preparation of HiC samples

Cells were harvested, and 1 M cells were fixed in 2% formaldehyde (Fisher Chemical) according to the ARIMA-HiC protocol. Samples were prepared and sequenced according to the manufacturer’s protocol by NYU Genome Technology Center.

#### Cellular Fractionation and Immunoprecipitation

All cellular fractionation and immunoprecipitation experiments were performed at 4° C or on ice with buffers containing 1 μg/ml Pepstatin, 1 μg/ml Aprotonin, 1 μg/ml Leupeptin, 0.3 mM PMSF, 10 mM Sodium fluoride, and 5 mM Sodium orthovanadate. For FLAG affinity purification from native chromatin (native ChIP-mass spectrometry), nuclear extracts in ESCs and MNs were prepared using Buffer A and Buffer C as described (*48*). Cytosolic fraction was removed by Buffer A (10 mM Tris, pH 7.9, 1.5 mM MgCl_2_, 10 mM KCl, and 0.5 mM dithiothreitol (DTT)). The pellet was resuspended in Buffer C (20 mM Tris, pH 7.9, 25% glycerol, 420 mM NaCl, 1.5 mM MgCl_2_, 0.2 mM EDTA, and 0.5 mM DTT), and incubated 1 hr to obtain nuclear extract. After removing nuclear extract, the remaining nuclear pellet was solubilized by benzonase (Millipore) digestion in a buffer containing 50 mM Tris, pH 7.9 and 2 mM MgCl_2_. For FLAG affinity purification from native chromatin and mass spectrometry, 20 mg nuclear pellet was incubated with 200 μl FLAG M2 beads in BC250 overnight and washed six times with BC250 containing 0.05% NP40, as described (*49*). Two elutions were performed with 0.5 mg/ml FLAG peptide in BC50 (without any protease inhibitors) with rotation at 4° C for 2 hr for a total of 4 hr. The eluate was sent for mass spectrometry to the Rutgers Mass Spectrometry Center and analyzed by liquid chromatography-mass spectrometry (LC-MS/MS). Peptide counts are shown for the native ChIP-mass spectrometry experiments in **Data S2**.

For FLAG affinity purification from crosslinked chromatin (crosslinked ChIP-mass spectrometry), a modified version of a previously reported protocol was applied (*31, 32*). Briefly, cells were crosslinked and sonicated as described above for ChIPseq except with a larger fragment size to include ~3-5 nucleosomes. 3 mg chromatin was used for FLAG affinity purification, and FLAG elutions were performed after stringent washes as described (*31*), but excluding the second step in the protocol wherein DNA is biotinylated. After de-crosslinking, samples were sent for mass spectrometry to the Rutgers Mass Spectrometry Center and analyzed by liquid chromatography-mass spectrometry (LC-MS/MS).

For extraction in 293FT cells, CβF expression vectors containing cDNAs for CTCF (mouse) or MAZ (mouse) were transfected into 293FT cells using PEI, and nuclei was prepared by using TMSD and BA450 buffers as described before (*50, 51*). Briefly, TMSD buffer (20 mM HEPES, 5 mM MgCl_2_, 85.5 g/L Sucrose, 25 mM NaCl, and 1 mM DTT) was used for cytosol removal, and nuclear extraction was done in BA450 buffer (20 mM HEPES, 450 mM NaCl, 5% glycerol, and 0.2 mM EDTA). FLAG affinity purification and FLAG peptide elution were performed similarly in the nuclear fraction.

#### Library Construction

All libraries were prepared according to the manufacturer’s instructions (Illumina). CRISPR libraries were prepared by performing two-step PCRs as described (*22*). Briefly, sgRNAs were amplified from genomic DNA by keeping the coverage maintained throughout the screens (300X for GeCKO v2 library, 1000X for custom library in secondary screens) and performing secondary amplifications by using Phusion polymerase (New England Biolabs) to attach Illumina adaptors (shown in **Table S3**). ChIPseq libraries were prepared as described (*16*). RNAseq libraries were prepared using KAPA library preparation kits, and HiC libraries were prepared according to the ARIMA standard HiC protocols by the NYU Genome Technology Center.

#### Data Analysis

##### CRISPR screens

MaGeCK tools were used for all primary and secondary CRISPR screen analysis (*26, 27*). Genome-wide screens with GeCKO v2 library A (3sgRNAs/gene) and GeCKO v2 library B (3sgRNAs/gene) were analyzed together in total populations of ESCs and MNs to identify essential/differentiation related genes (negative selections). The analysis was done separately for library A (2 screens) and library B (2 screens) in sorted MN populations to identify genes affecting CTCF boundary function (positive selection). PANTHER database was used for Gene Ontology (GO) analysis (*52*). To generate Venn diagrams in CRISPR screens, web-tools at http://genevenn.sourceforge.net were used.

#### RNAseq

RNA-seq data was analyzed as described (*16*). Briefly, sequence reads were mapped to mm10 reference genome with Bowtie 2 (*53*) and normalized differential gene expression was obtained with DEseq (R package) (*54*). Relevant expression and p-values are listed for differentially expressed genes in **Data S7-S10**. PANTHER database was used for Gene Ontology (GO) analysis (*52*).

#### ChIPseq

ChIPseq experiments were analyzed as before (*47*). In brief, sequence reads were mapped to mm10 reference genome with Bowtie 2 using default parameters (*53*). After normalization with the spike-in *Drosophila* read counts, normalized ChIP-seq read densities were visualized in Integrative Genomics Viewer (IGV) (*55*). Heat maps were generated using deepTools in R (*56*). ‘ChIPpeakAnno’ package from Bioconductor (*57*) was used to draw Venn diagrams to visualize the overlap among ChIP-seq samples. The replicates were assessed similarly by visualizing at IGV and generating heat maps. ChIPseq “bed” file coordinates were converted into “fasta” by using fetch sequences tool within Regulatory Sequence Analysis Tools (RSAT) (*58*), and MEME was used for motif analysis of MAZ in ESCs and MNs (*59*).

#### HiC

All samples were prepared in two biological replicates. All Hi-C data were analyzed by the hic-bench platform (*60*). Throughout our comprehensive analysis, the following operations were done using hic-bench. Internally, bowtie2 (*61*) was used to align the paired reads using mm10 reference genome and only the read pairs uniquely mapped to the same chromosome with the mapping quality ≥ 20 and the pair distance ≥ 25 kb were used. Then, the interaction matrix was tabulated by reading the coordinates of aligned reads in 20 kilobase bins. To ensure that each interaction bin showed equal visibility, the iterative correction method (*62*) was used to normalize the bins. For the compartment analysis, the Hi-C interaction bins were divided into A and B compartments using the first principal component values from HOMER’s runHiCpca (*63, 64*). Using hic-bench, the compartment changes from comparison of two cell types for the bins in the interaction matrix were visualized by the stacked bar plot.

Topologically associated domains (TADs) were defined as shown (*60, 65*) with the insulating window of 500 kb. The boundaries of TADs were called from the boundary score, using the “ratio” method defined (*60*), wherein each TAD boundary had a noticeably lower boundary score than the neighboring region. The score was calculated for each 20 kilobase bin using the window size of 250 kb, 500 kb, and 1000 kb. In the principal component analysis to distinguish the differences, the boundary score for every replicate and cell type was combined, quantile normalized, and plotted. Then, for each TAD, the magnitude of intra-TAD “activity” was defined as in (*65*). The cutoff for significantly differential TADs was Benjamini-Hochberg corrected Q value of 0.05 and no cutoff for the fold change.

Significantly enriched chromatin loops were called using FitHi-C 2 (*66*) with default parameters. To characterize the loops by the level of CTCF and MAZ ChIP-seq level, aggregate peak analysis (APA) software (*67*) was used to show the averaged profile. The genome sequence that matched the transcription factor motifs of mouse CTCF and MAZ from the Catalog of Inferred Sequence Binding Preferences (CIS-BP) (*68*) was found from PWMScan (*69*). Visualization of Hi-C and associated ChIP-seq data was made with pyGenomeTracks (*70*).

**Fig. S1.**
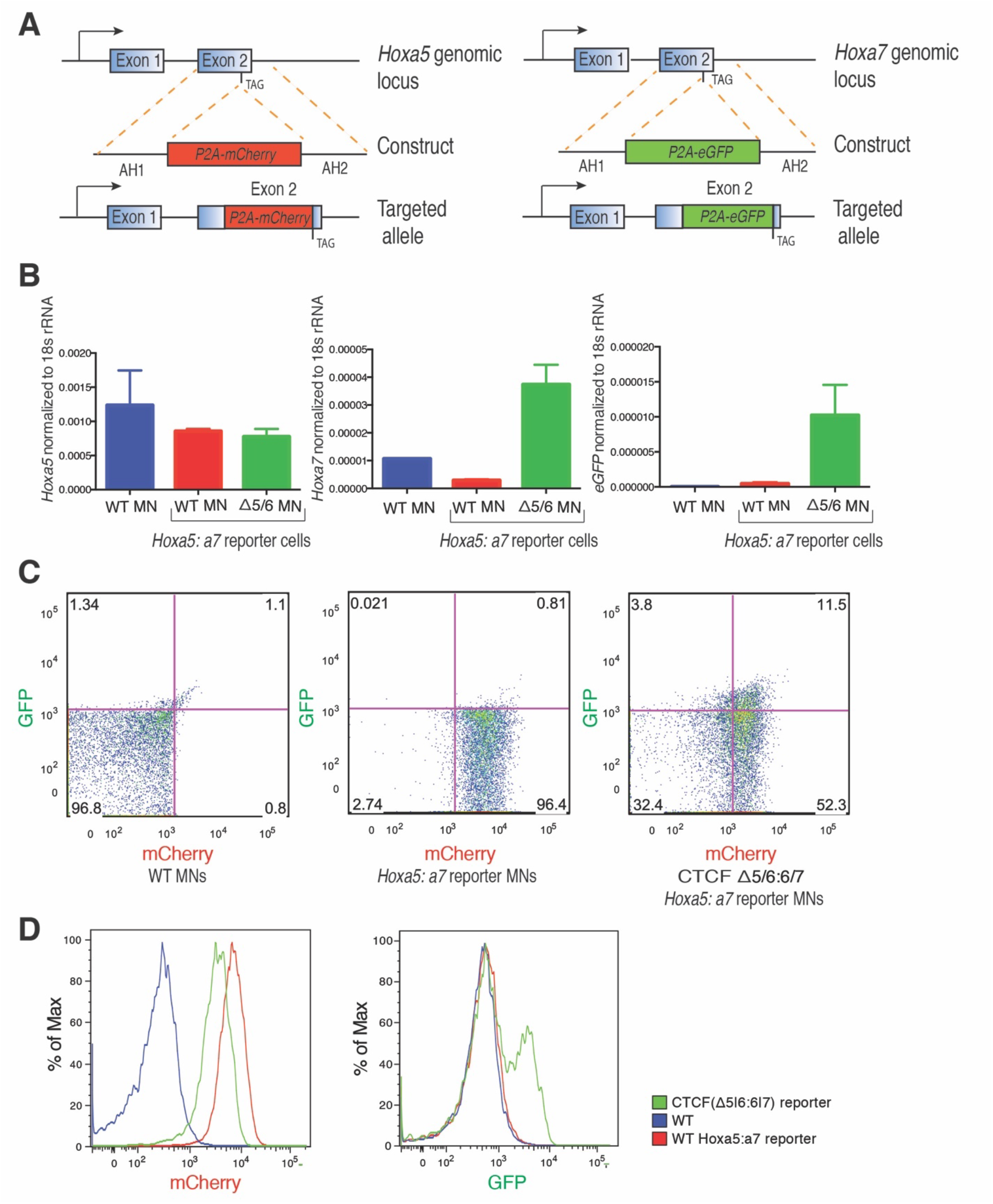
A dual-fluorescent reporter *Hoxa5:a7* mESC line can be used to assess disruption of a functional CTCF boundary in MNs. **A.** Strategy for generating the *Hoxa5:a7* reporter mESC line via CRISPR. AH1, AH2: arm of homology 1, 2. **B.** RT-qPCR signal for *Hoxa5, Hoxa7,* and *eGFP* normalized to 18s ribosomal RNA in WT MNs, *Hoxa5:a7* reporter MNs, and CTCF(Δ5|6) *Hoxa5:a7* reporter MNs. Error bars indicate the standard deviation across two biological replicates. MNs were obtained through *in vitro* differentiation of mESCs. **C.** FACS data showing mCherry and eGFP reporter expression in WT MNs (left), *Hoxa5:a7* reporter MNs (middle), and CTCF(Δ5|6:6|7) *Hoxa5:a7* reporter MNs (right). These plots are one representative of three biological replicates. Percentage of positive population in each quadrant is indicated. **D.** Flow cytometry analysis of mCherry and eGFP reporter expression in MNs with the indicated genotypes: WT, *Hoxa5:a7* reporter, and CTCF(Δ5|6:Δ6|7) reporter (see **Fig. S6** for RT-qPCR of *Hox* genes). These plots are one representative of three biological replicates. Δ5|6 refers to CTCF binding site deletion between *Hoxa5*-*6.* Δ5|6:6|7 stands for 2 CTCF binding site deletions.

**Fig. S2.**
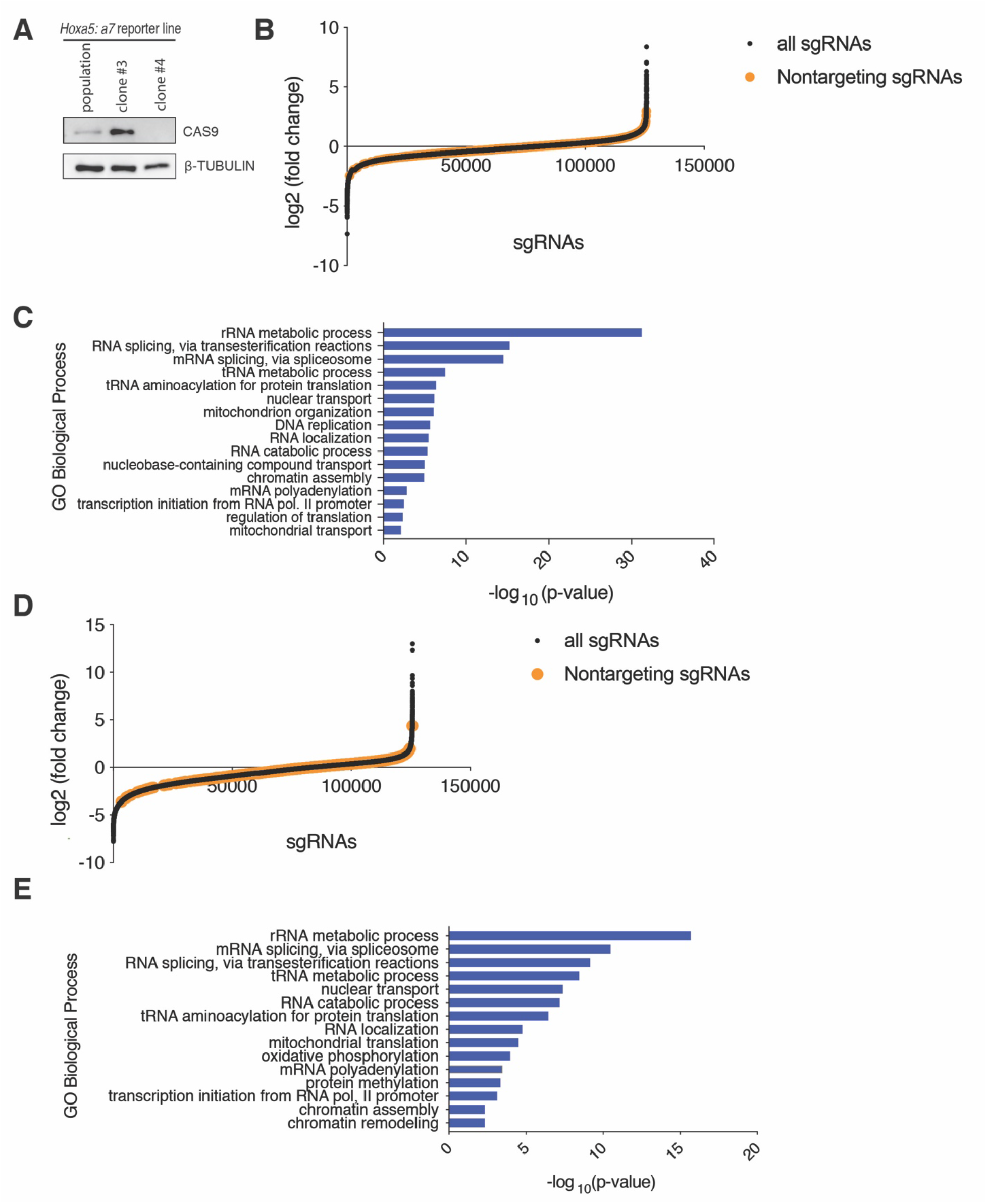
Genome-wide CRISPR KO screen shows selective loss of essential genes in ESCs, and MNs. **A.** CAS9 protein expression in dual reporter mESCs transduced with *Cas9* lentivirus and selected with blasticidin. β-TUBULIN acts as a loading control. **B.** Fold change of sgRNAs in the primary screens comparing ESCs to plasmid library. **C.** Gene Ontology (GO) analysis of biological processes in negatively selected genes in ESCs compared to plasmid library (FDR<0.05). **D.** Fold change of sgRNAs in the primary screens comparing MN population to ESC population. **E.** GO analysis of biological processes in negatively selected genes in MNs compared to ESCs (FDR<0.05).

**Fig. S3.**
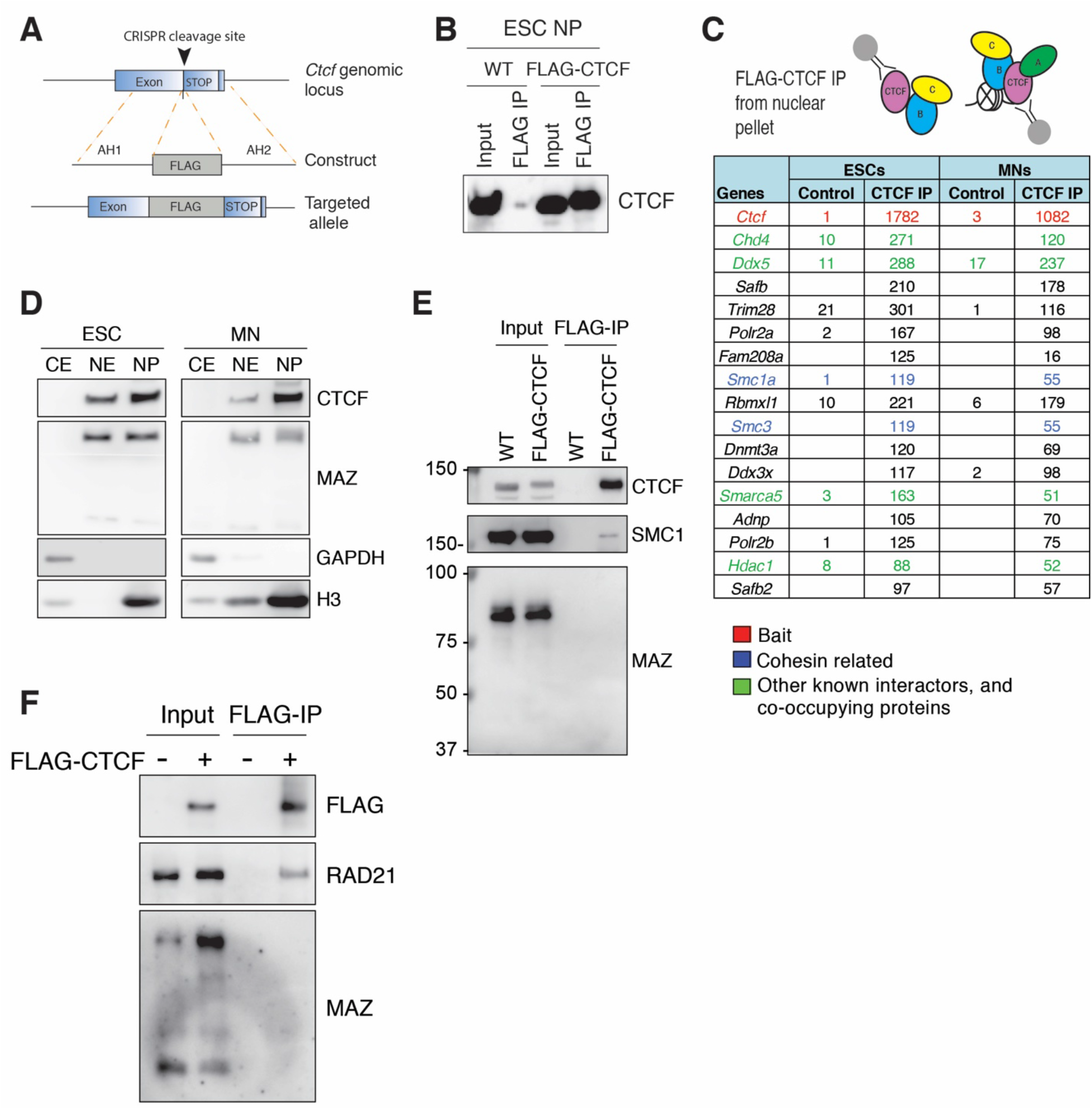
Identification of proteins co-localizing with CTCF via native ChIP-MS in ESCs and MNs. **A.** FLAG-tag integrated at the C-terminus of CTCF via CRISPR genome editing. AH1, 2: arm of homology 1, 2. **B**. FLAG pull-down followed by CTCF western blot in benzonase solubilized nuclear pellet (NP) of ESCs. **C**. Native FLAG-CTCF IP in ESCs and MNs results in identification of known CTCF interactors, and novel proteins. The mean peptide counts from two biological replicates of FLAG-CTCF IPs were normalized to the control FLAG IP from untagged cells. Candidates filtered from the top of the list are shown (see **Data S2** for all). **D.** Western blot analysis of CTCF and MAZ in different cellular fractions in mESCs and MNs. CE: cytoplasmic extract, NE: nuclear extract, NP: nuclear pellet. **E.** Western blot analysis of CTCF, SMC1, and MAZ upon FLAG-CTCF immunoprecipitation from nuclear pellet of mESCs. **F.** Western blot analysis of FLAG, RAD21, and MAZ upon FLAG-CTCF immunoprecipitation from nuclear extract of 293FT cells.

**Fig. S4.**
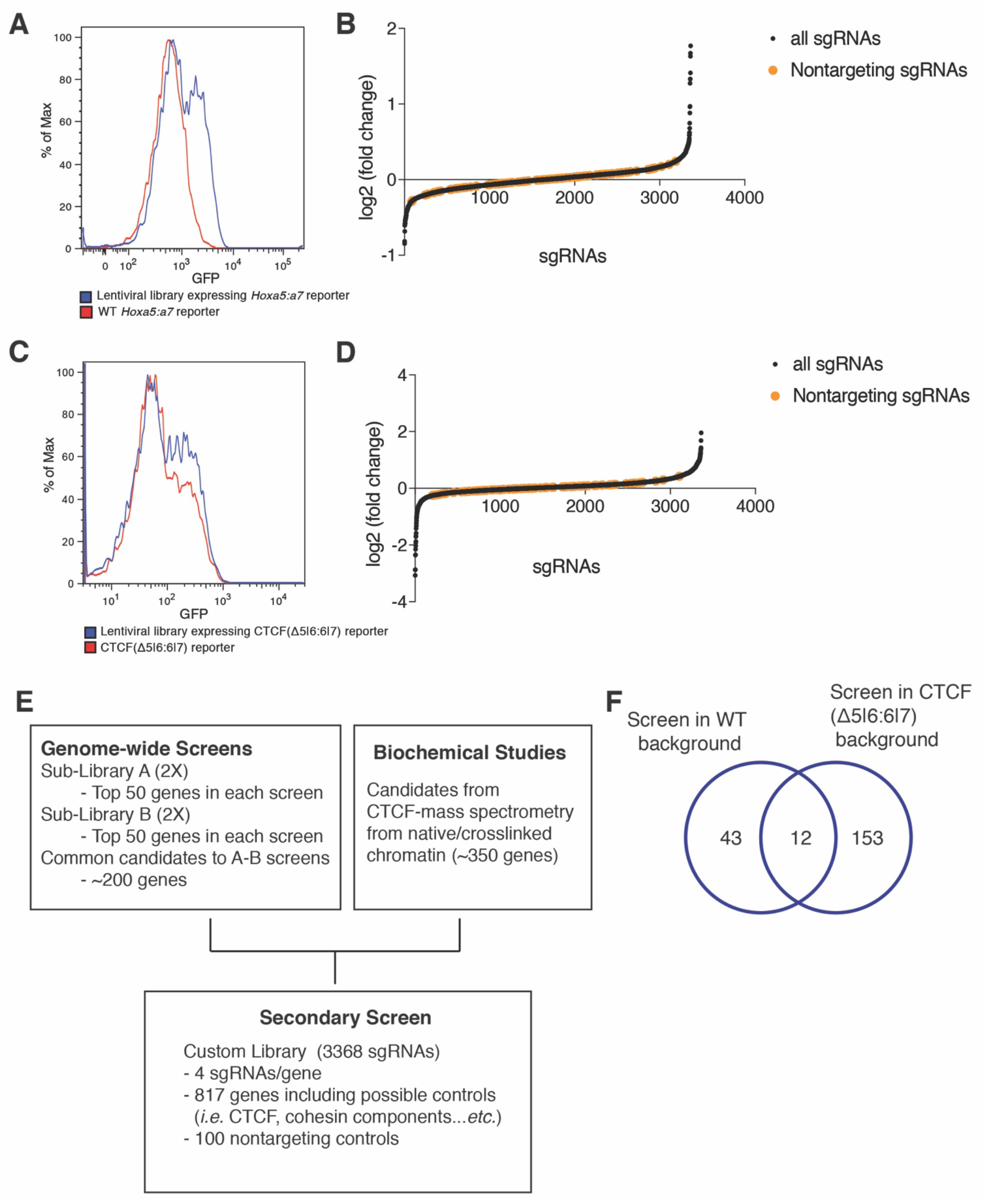
Secondary loss-of-function screen at the CTCF boundary in the *HoxA* cluster results in a small list of candidates. **A.** FACS analysis of GFP expression in lentivirus library expressing MNs in WT background (with intact CTCF binding sites) versus untransduced WT MNs. **B.** Fold change of sgRNAs in the secondary screen performed in WT background. **C.** FACS analysis of GFP expression in lentivirus library expressing MNs in CTCF(Δ5|6:6|7) background versus untransduced CTCF(Δ5|6:6|7) MNs. **D.** Fold change of sgRNAs in the secondary screen performed in CTCF(Δ5|6:6|7) background. **E.** Schematic of candidate selection for the secondary loss-of-function screens. **F.** Venn diagram depicting overlap of secondary genetic screens in WT versus CTCF(Δ5|6:6|7) background.

**Fig. S5.**
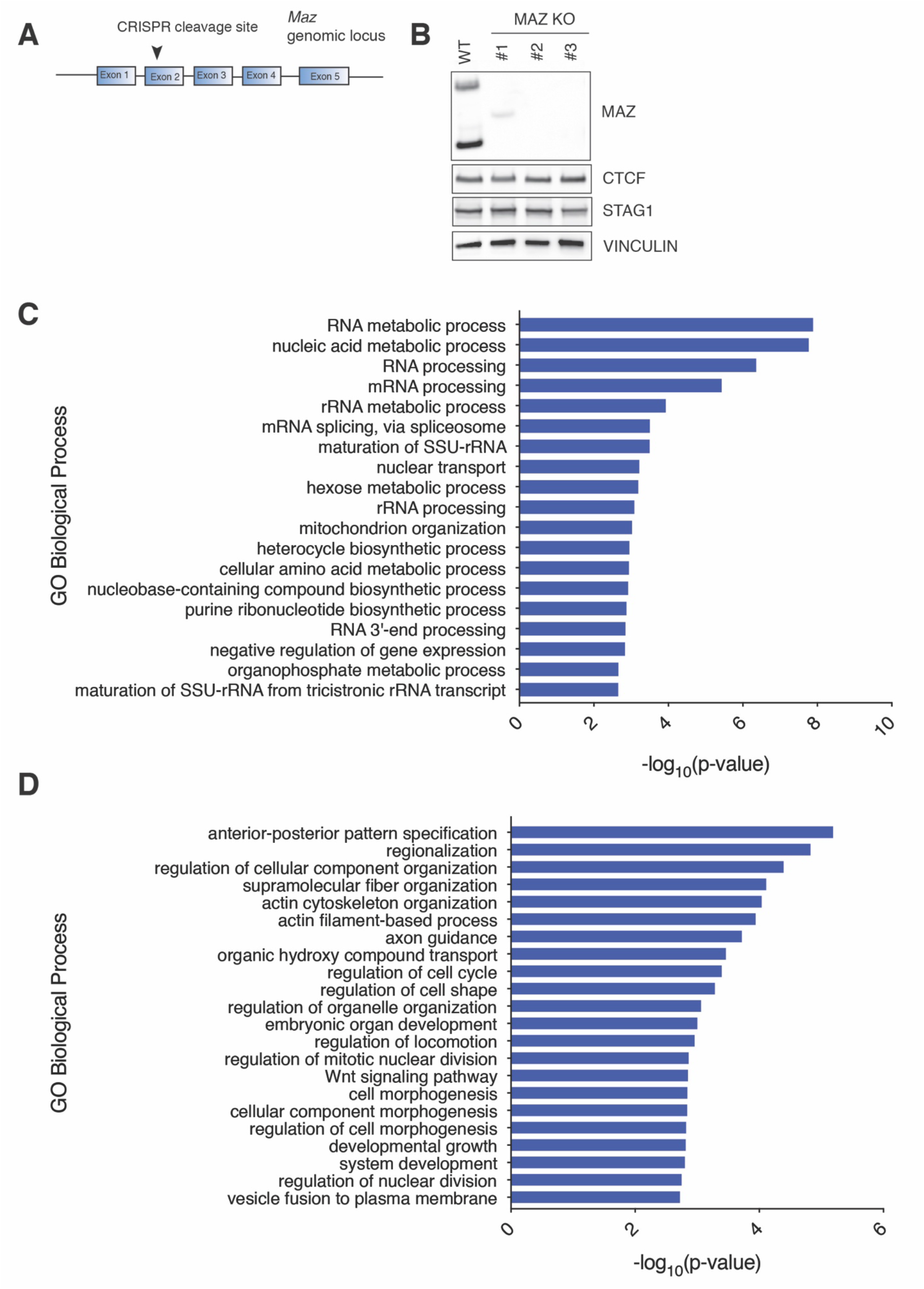
Developmental and neuronal processes are enriched upon loss of *Maz* in motor neurons. **A**. The strategy of MAZ KO mESC line generation via CRISPR. **B**. Western blot analysis of indicated proteins in WT and MAZ KO ESCs. **C.** GO analysis of biological processes in differentially expressed genes in MAZ KO ESCs compared to WT ESCs. **D.** GO analysis of biological processes in differentially expressed genes in MAZ KO MNs compared to WT MNs.

**Fig. S6.**
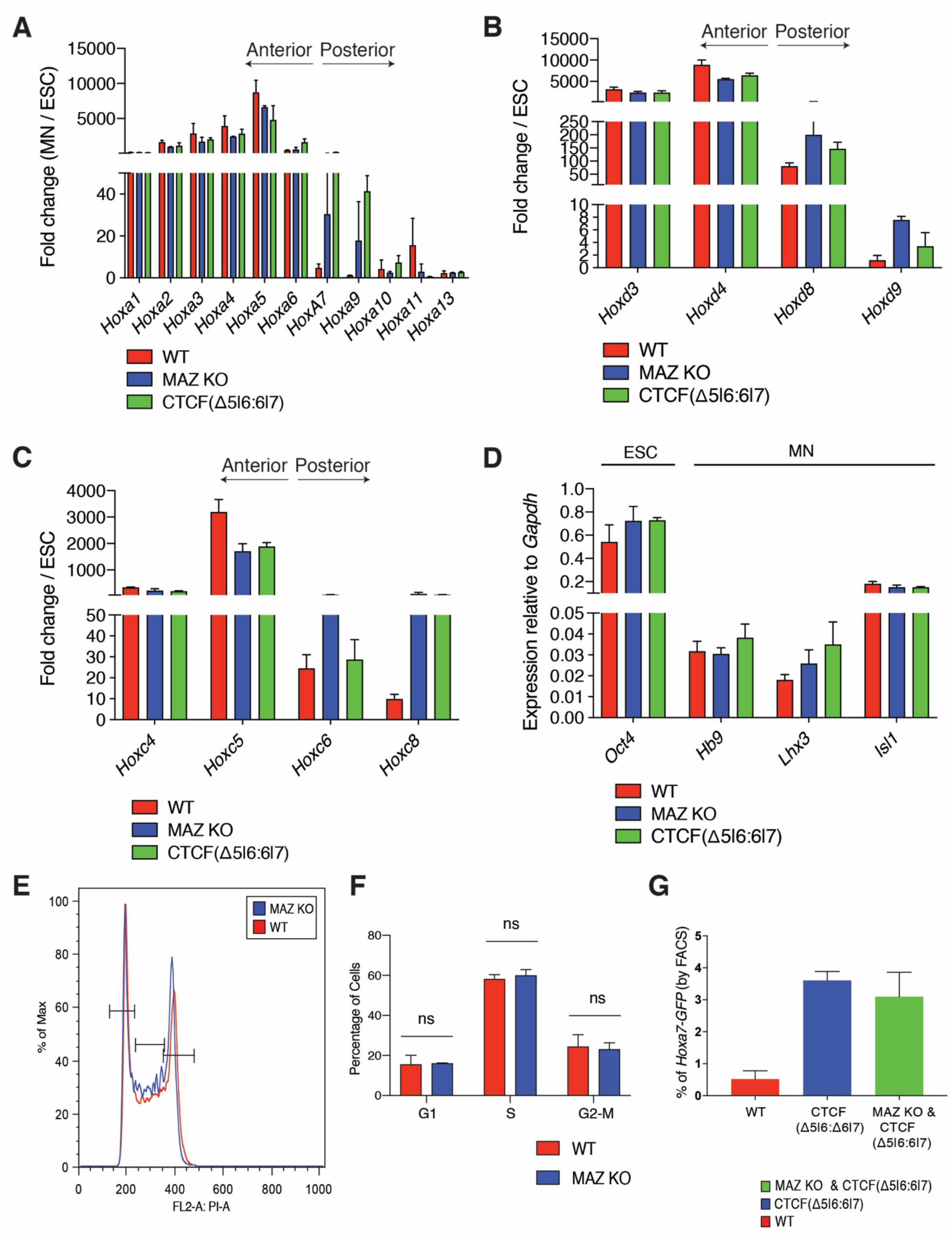
Loss of *Maz* results in *Hox* gene de-repression downstream of CTCF boundaries at *HoxA*, *HoxD*, and *HoxC* clusters. **A.** RT-qPCR analysis for the indicated *Hox* genes in *HoxA* cluster, *HoxD* cluster (**B**), and *HoxC* cluster (**C**) in WT, MAZ KO, and CTCF(Δ5|6:6|7) cells. RT-qPCR signal is normalized to *Gapdh* levels. Fold change in expression in MNs is calculated relative to baseline expression in ESCs. **D.** RT-qPCR analysis for the indicated ESC and MN markers in WT, MAZ KO, and CTCF(Δ5|6:6|7) cells. RT-qPCR signal is normalized to *Gapdh* levels. Error bars in all RT-qPCR results represent standard deviation across three biological replicates. Maz KO represents three independent clones. **E.** FACS analysis of cell cycle in WT versus MAZ KO ESCs. **F.** Quantification of cell cycle analysis by FACS in WT versus MAZ KO ESCs. Error bars indicate standard deviation across three biological replicates. Maz KO represents three independent clones. **G**. The percentage of *Hoxa7-GFP* cells quantified based on FACS analysis in MNs with the indicated genotypes: WT, CTCF(Δ5|6:6|7), and MAZ KO & CTCF(Δ5|6:6|7). Error bars show standard deviation across three biological replicates. Results from MAZ KO & CTCF(Δ5|6:6|7) represent three independent clones.

**Fig. S7.**
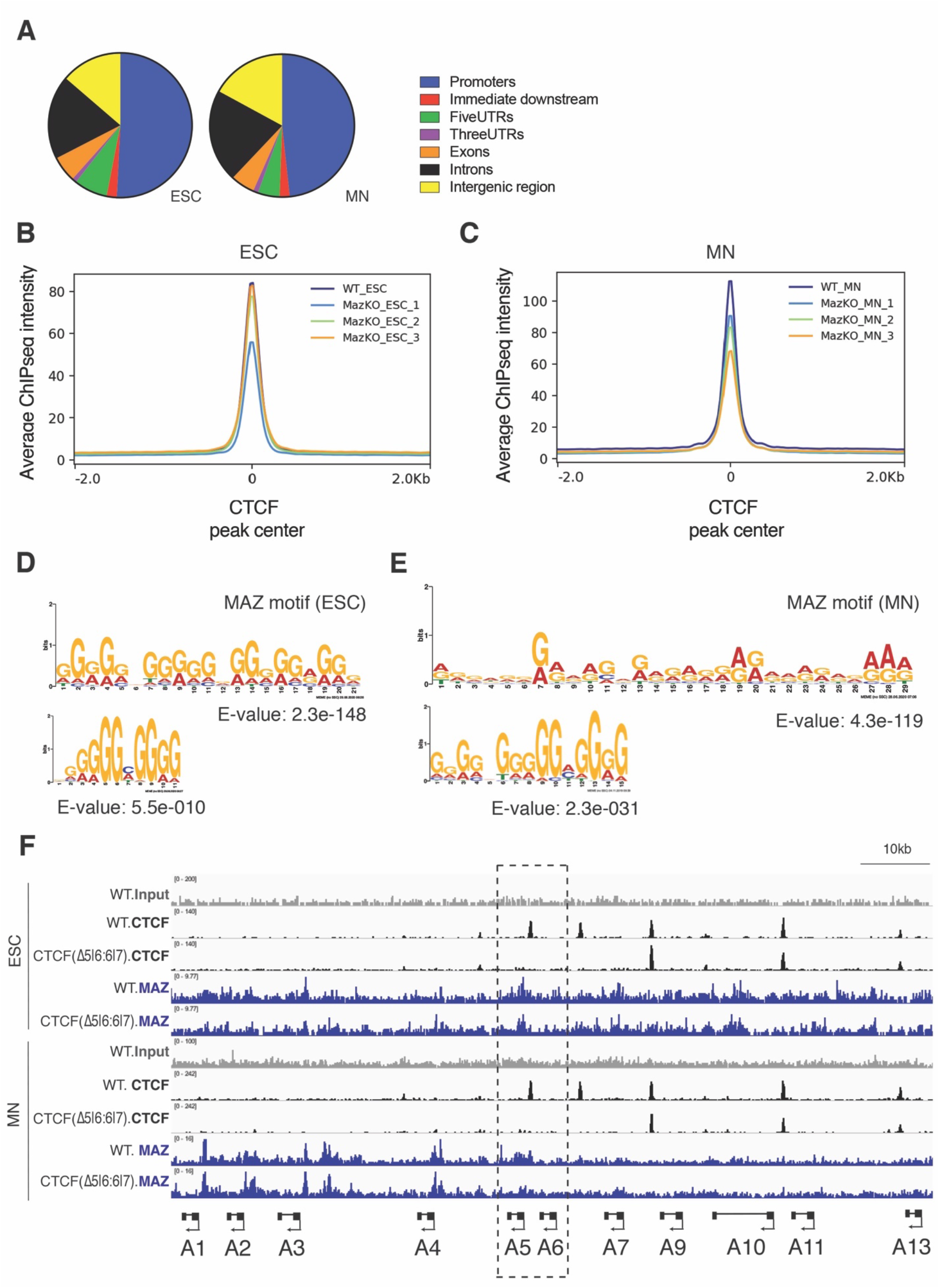
CTCF binding is altered globally upon MAZ KO in ESCs and MNs. **A.** Distribution of MAZ binding sites across genomic features. **B**. Average CTCF ChIPseq intensity plotted for WT versus MAZ KO ESCs and MNs (**C**) globally within a 4 kb window centered at CTCF peak center from three biological replicates. **D.** Motif analysis of MAZ ChIPseq in ESCs, and MNs (**E**) by using MEME. Top two motifs are shown. **F.** Normalized ChIP-seq densities for CTCF, and MAZ in WT, and CTCF (Δ5|6:6|7) ESCs and MNs in the *HoxA* cluster. ChIPseq tracks are from one representative of two biological replicates for CTCF and MAZ except for one replicate for MAZ in CTCF (Δ5|6:6|7) ESCs and MNs.

**Fig. S8.**
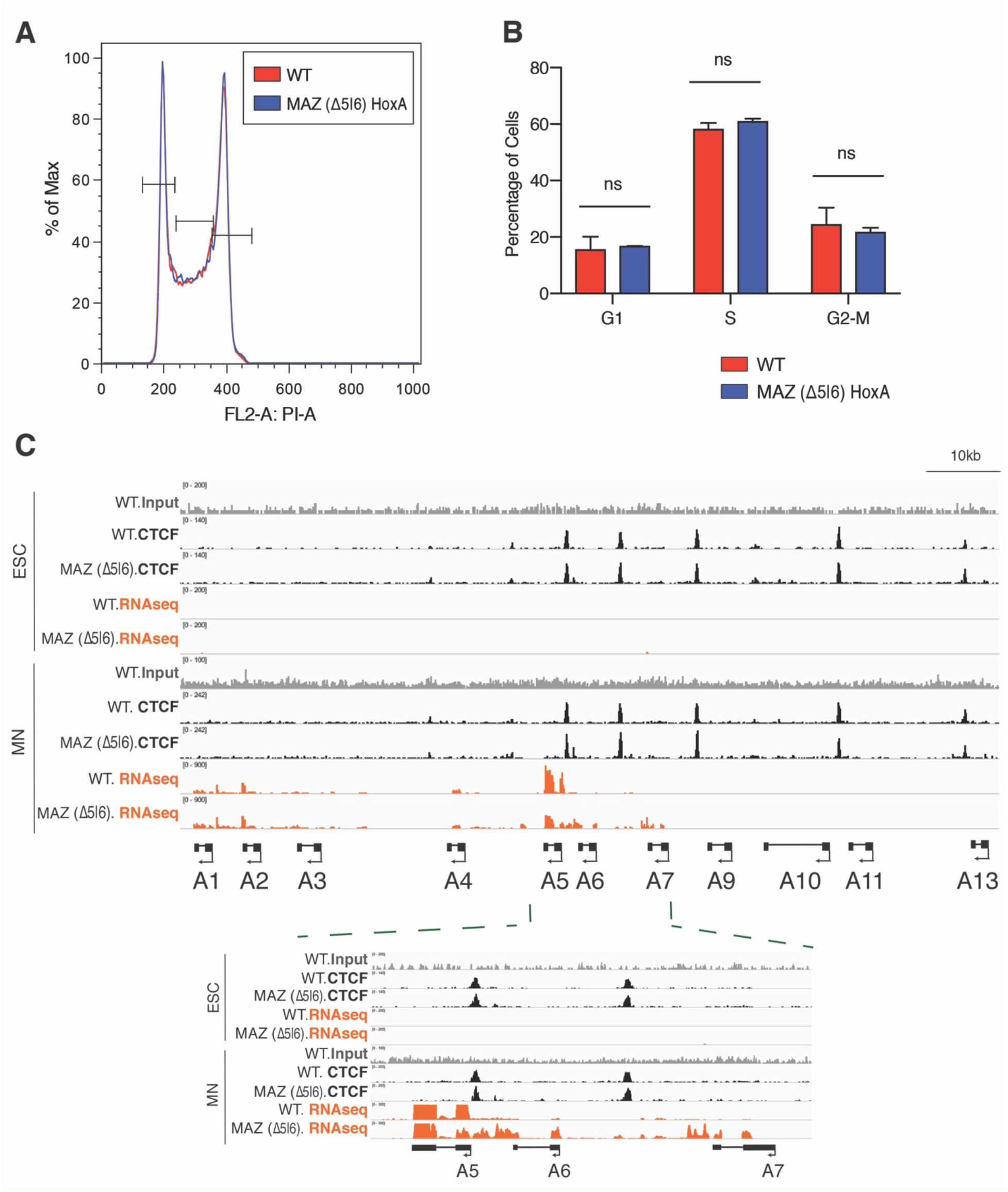
Loss of MAZ binding site at *HoxA* cluster alters *Hox* gene expression pattern at *HoxA* cluster. **A.** FACS analysis of cell cycle in WT versus MAZ (Δ5|6) ESCs. **B.** Quantification of cell cycle analysis by FACS in WT versus MAZ (Δ5|6) ESCs. Error bars represent standard deviation across three biological replicates. Student’s t-test was used. **C.** Normalized ChIP-seq densities for CTCF, and RNAseq tracks in WT, and MAZ (Δ5|6) ESCs and MNs in the *HoxA* cluster. Below is an enlarged view. ChIPseq tracks are from one representative of two biological replicates for CTCF. RNAseq tracks are from one representative of three biological replicates.

**Fig. S9.**
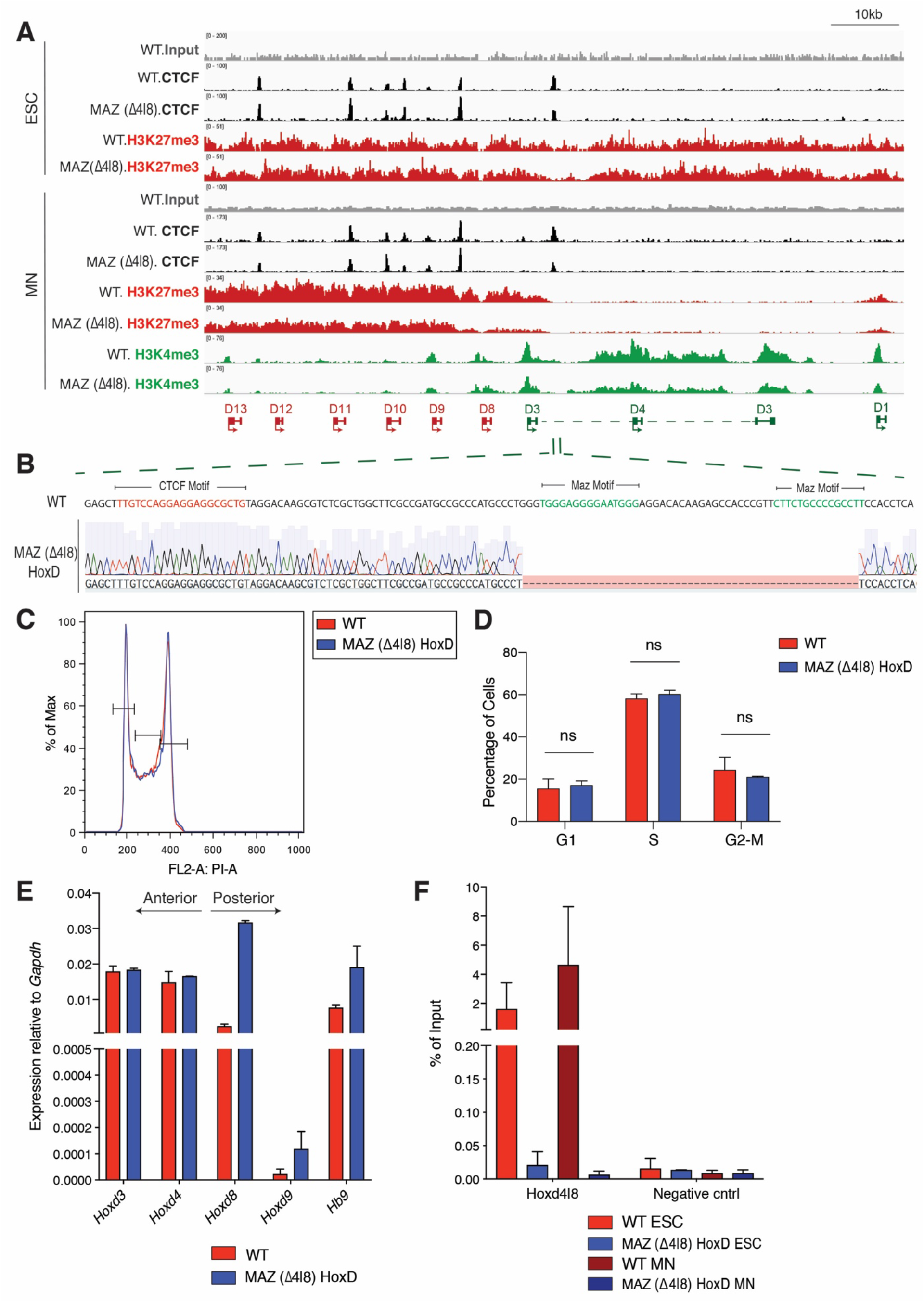
Loss of MAZ binding site alters *Hox* gene expression pattern and chromatin domains at *HoxD* cluster. **A**. Normalized ChIP-seq densities for CTCF and indicated histone post-translational modifications (PTMs) in WT, and MAZ (Δ4|8) ESCs and MNs in the *HoxD* cluster. ChIPseq tracks are from one representative of two biological replicates for CTCF, and one replicate for histone PTMs. **B**. MAZ binding site deletion via CRISPR is depicted for the 4|8 site at the *HoxD* cluster. **C.** FACS analysis of cell cycle in WT versus MAZ (Δ4|8) ESCs. **D.** Quantification of cell cycle analysis by FACS in WT versus MAZ (Δ4|8) ESCs. Error bars represent standard deviation across three biological replicates. Student’s t-test was used. **E**. RT-qPCR for the indicated *Hox* genes in the *HoxD* cluster in MNs upon MAZ (Δ4|8). Error bars indicate standard deviation across two biological replicates. **F**. MAZ ChIP-qPCR analysis in the *HoxD* cluster in mESCs and MNs upon MAZ (Δ4|8). Error bars show standard deviation across two biological replicates.

**Fig. S10.**
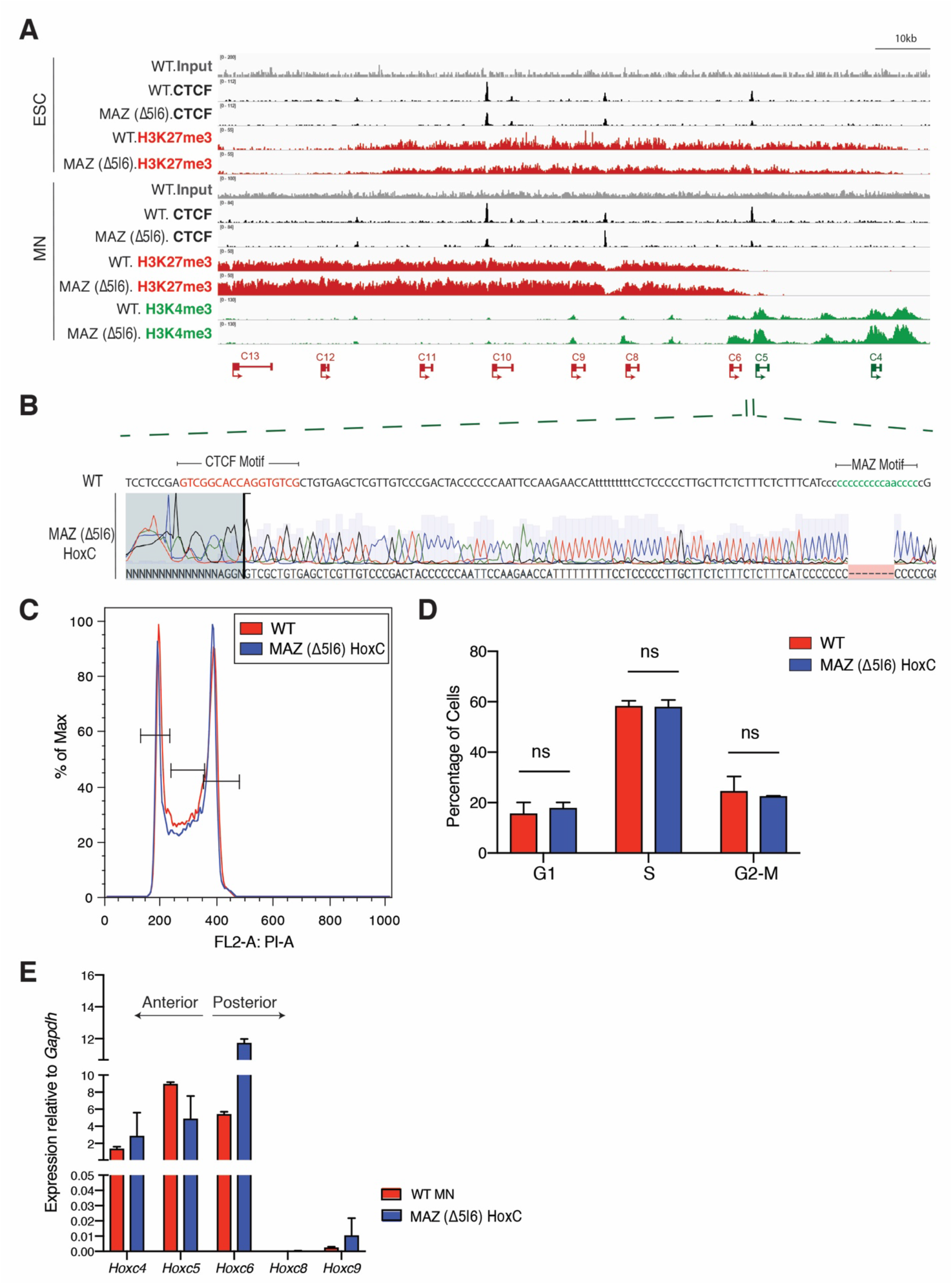
Loss of MAZ binding site alters *Hox* gene expression pattern and chromatin domains at the *HoxC* cluster. **A**. Normalized ChIP-seq densities for CTCF, and indicated histone PTMs in WT, and MAZ (Δ5|6) ESCs and MNs in the *HoxC* cluster. ChIPseq tracks represent one biological replicate. **B**. MAZ binding site deletion via CRISPR is depicted for the 5|6 site at the *HoxC* cluster. **C.** FACS analysis of cell cycle in WT versus MAZ (Δ5|6) ESCs. **D.** Quantification of cell cycle analysis by FACS in WT versus MAZ (Δ5|6) ESCs. Error bars represent standard deviation across three biological replicates. Student’s t-test was used. **E**. RT-qPCR for the indicated *Hox* genes in the *HoxC* cluster in MNs upon MAZ (Δ5|6). Error bars show standard deviation across two biological replicates.

**Fig. S11.**
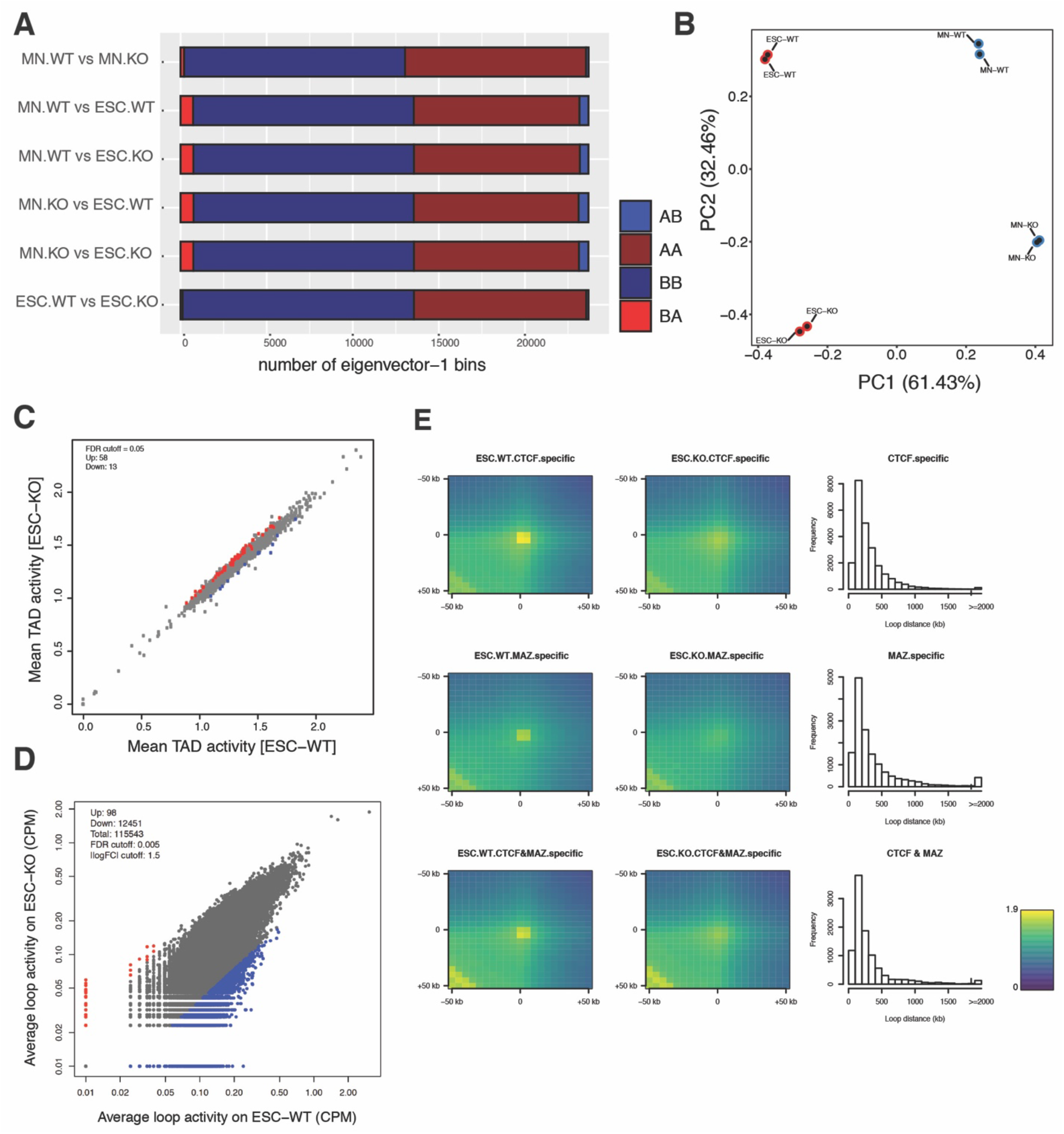
HiC analysis of global organization in MAZ KO ESCs and MNs. **A.** Bar plot of compartment switches between active (A) and inactive (B) compartments in ESCs and MNs. **B.** Principal Component Analysis (PCA) of boundary scores in WT ESCs, MAZ KO ESCs, WT MNs, and MAZ KO MNs. **C.** Scatter plot showing differential intra-TAD activity in WT vs MAZ KO ESCs (FDR cut-off=0.05). **D.** Scatter plot showing differential loop activity in WT vs MAZ KO ESCs (all loops, n=115543, FDR cut-off=0.005, | log (Fold Change) | cut-off=1.5, Up-regulated=98, Down-regulated: 12451). **E.** Aggregate Peak Analysis (APA) of loops in WT vs MAZ KO ESCs showing ChIP-seq signals of CTCF, MAZ, or both at any region covered by them. The resolution of APA is 5 kb. Histograms showing the distribution of loop distance in MAZ KO compared to WT related to the binding level of ChIP-seq.

**Fig. S12.**
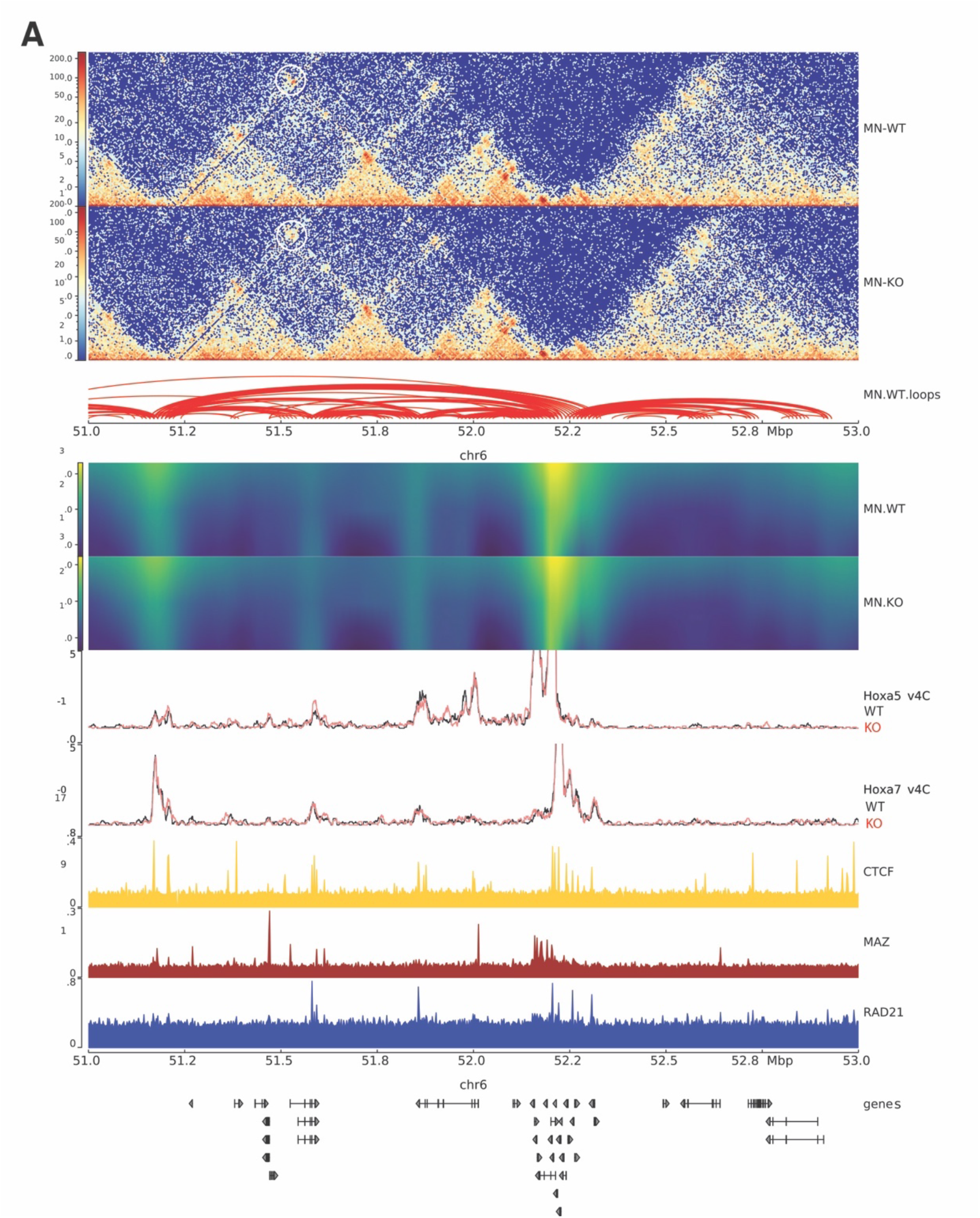
Visualization of HiC analysis at the indicated region in WT and MAZ KO MNs. **A.** Visualization of Hi-C contact matrices for a zoomed-in region on chromosome 6 with a change of loop activity in WT vs MAZ KO MNs. Shown below are loops identified in WT MNs, insulation score heat maps in WT MNs and MAZ KO MNs, virtual 4C plots with *Hoxa5* as viewpoint and *Hoxa7* as viewpoint, and ChIPseq read densities for CTCF, MAZ, and RAD21. The last track shows gene annotations.

**Fig. S13.**
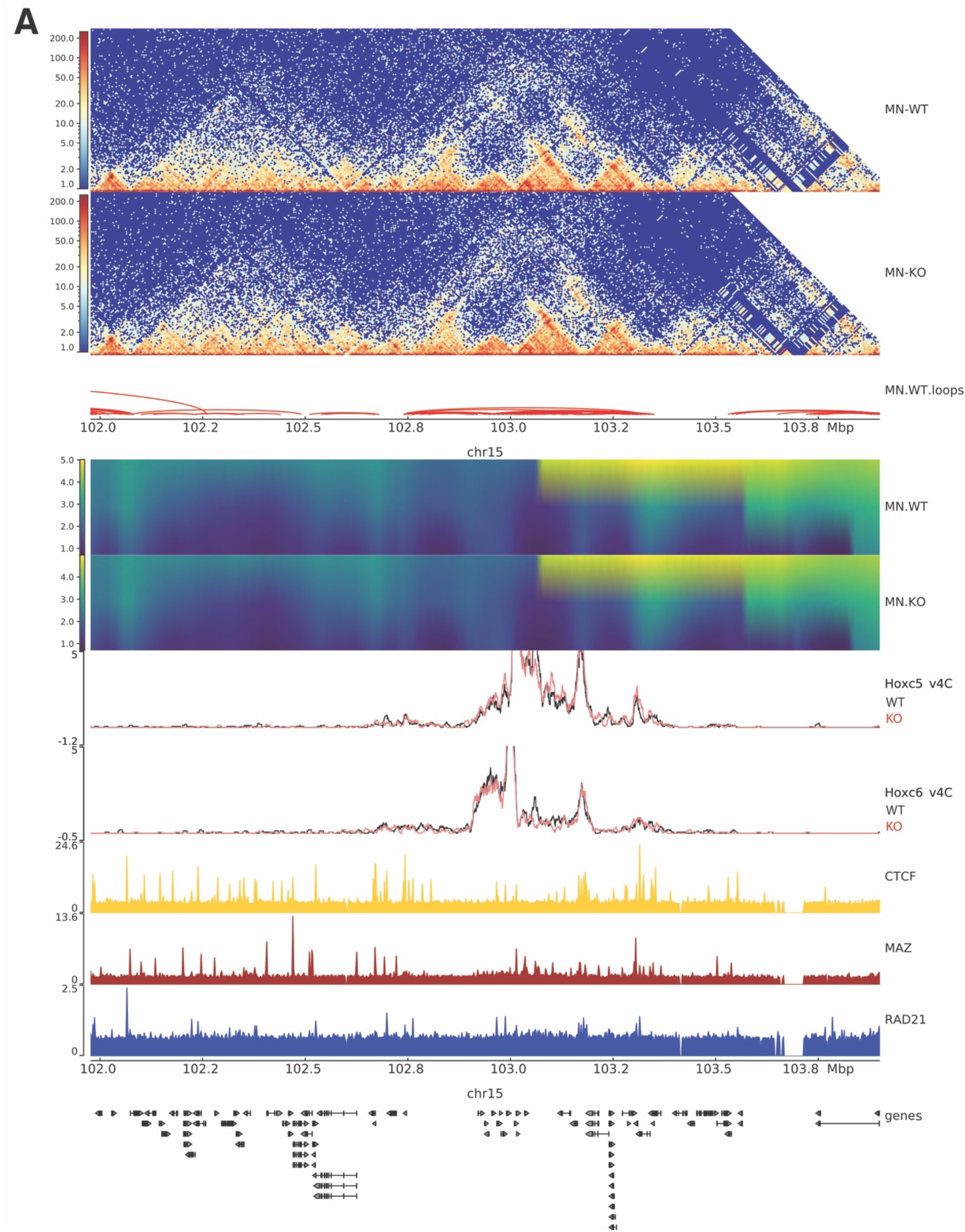
Visualization of HiC analysis at the indicated region in WT and MAZ KO MNs. **A.** Visualization of Hi-C contact matrices for a zoomed-in region on chromosome 15 in WT vs MAZ KO MNs. Shown below are loops identified in WT MNs, insulation score heat maps in WT MNs and MAZ KO MNs, virtual 4C plots with *Hoxc5* as viewpoint and *Hoxc6* as viewpoint, and ChIPseq read densities for CTCF, MAZ, and RAD21. The last track shows gene annotations.

**Fig. S14.**
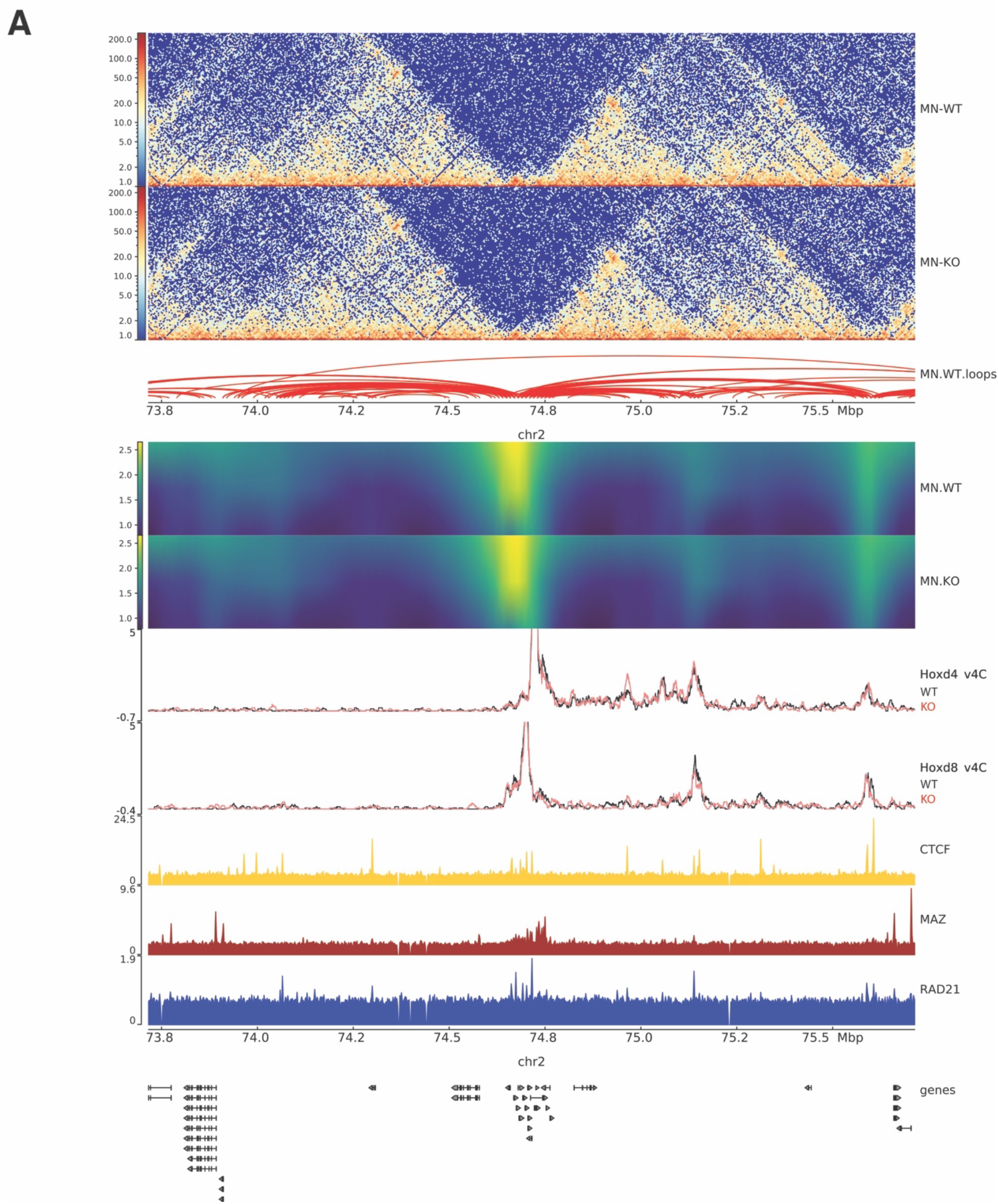
Visualization of HiC analysis at the indicated region in WT and MAZ KO MNs. **A.** Visualization of Hi-C contact matrices for a zoomed-in region on chromosome 2 in WT vs MAZ KO MNs. Shown below are loops identified in WT MNs, insulation score heat maps in WT MNs and MAZ KO MNs, virtual 4C plots with *Hoxd4* as viewpoint and *Hoxd8* as viewpoint, and ChIPseq read densities for CTCF, MAZ, and RAD21. The last track shows gene annotations.

**Table S1.**
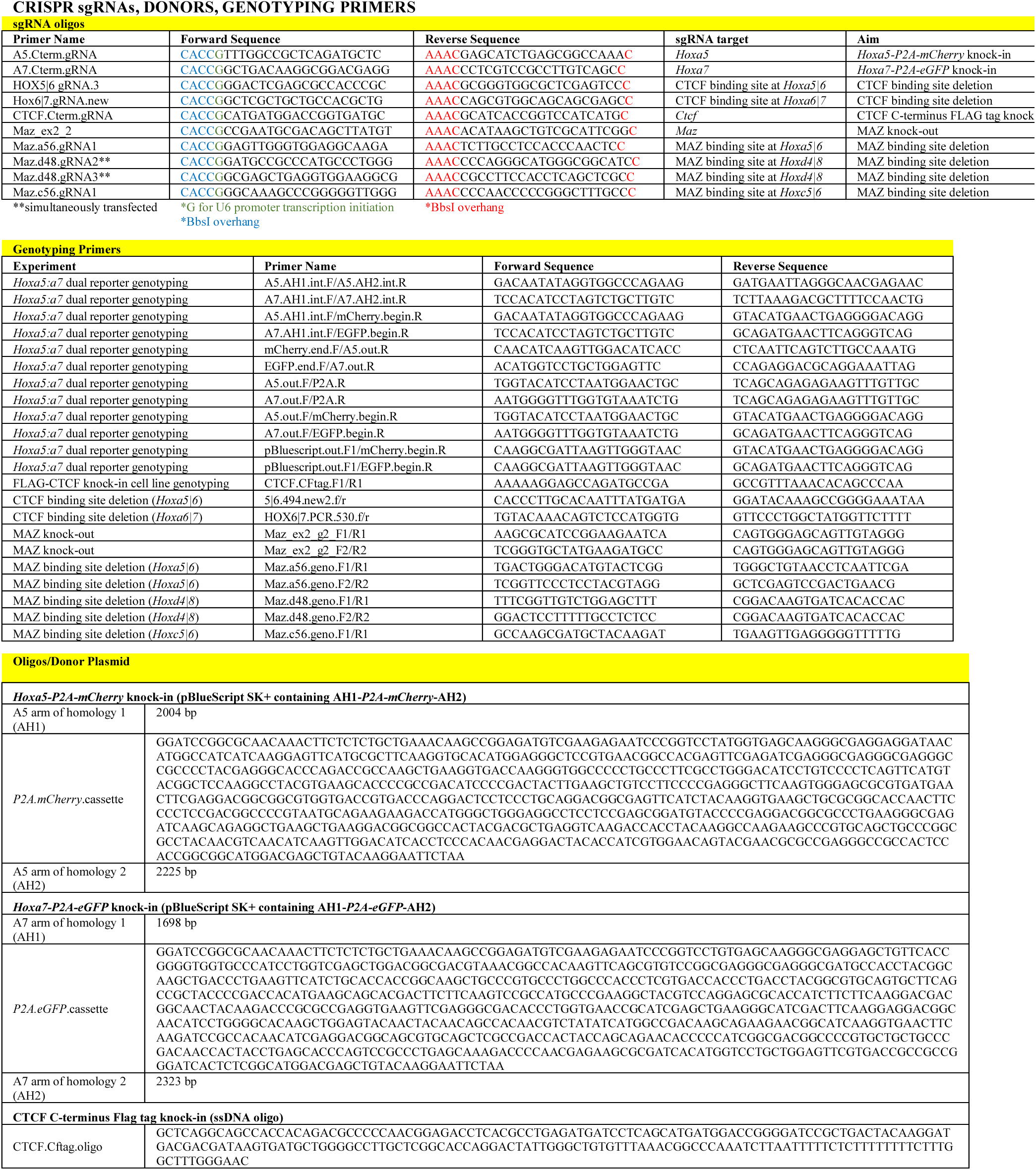
List of CRISPR sgRNAs, donors, and genotyping primers.

**Table S2.**
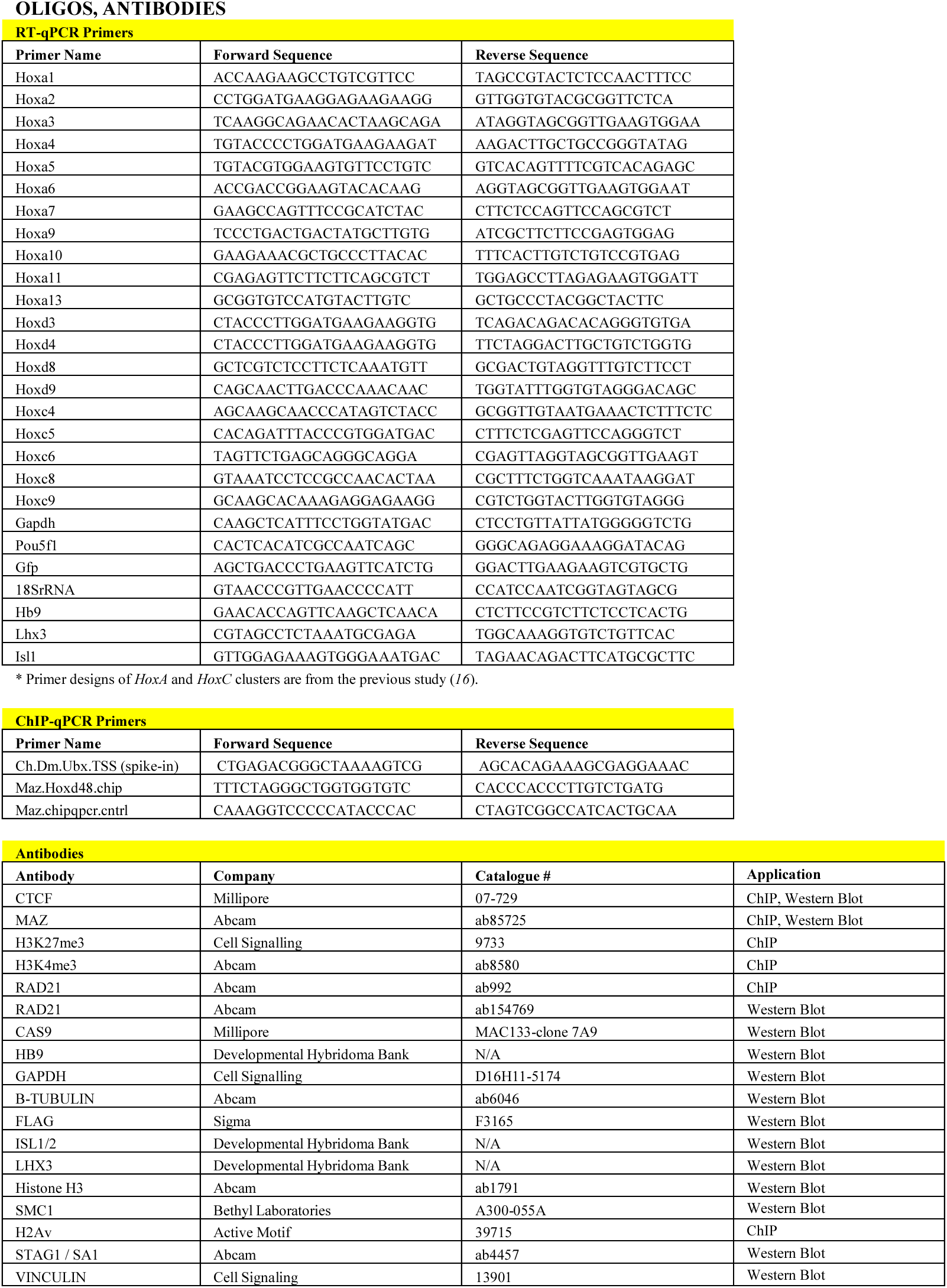
List of oligos and antibodies.

**Table S3.**
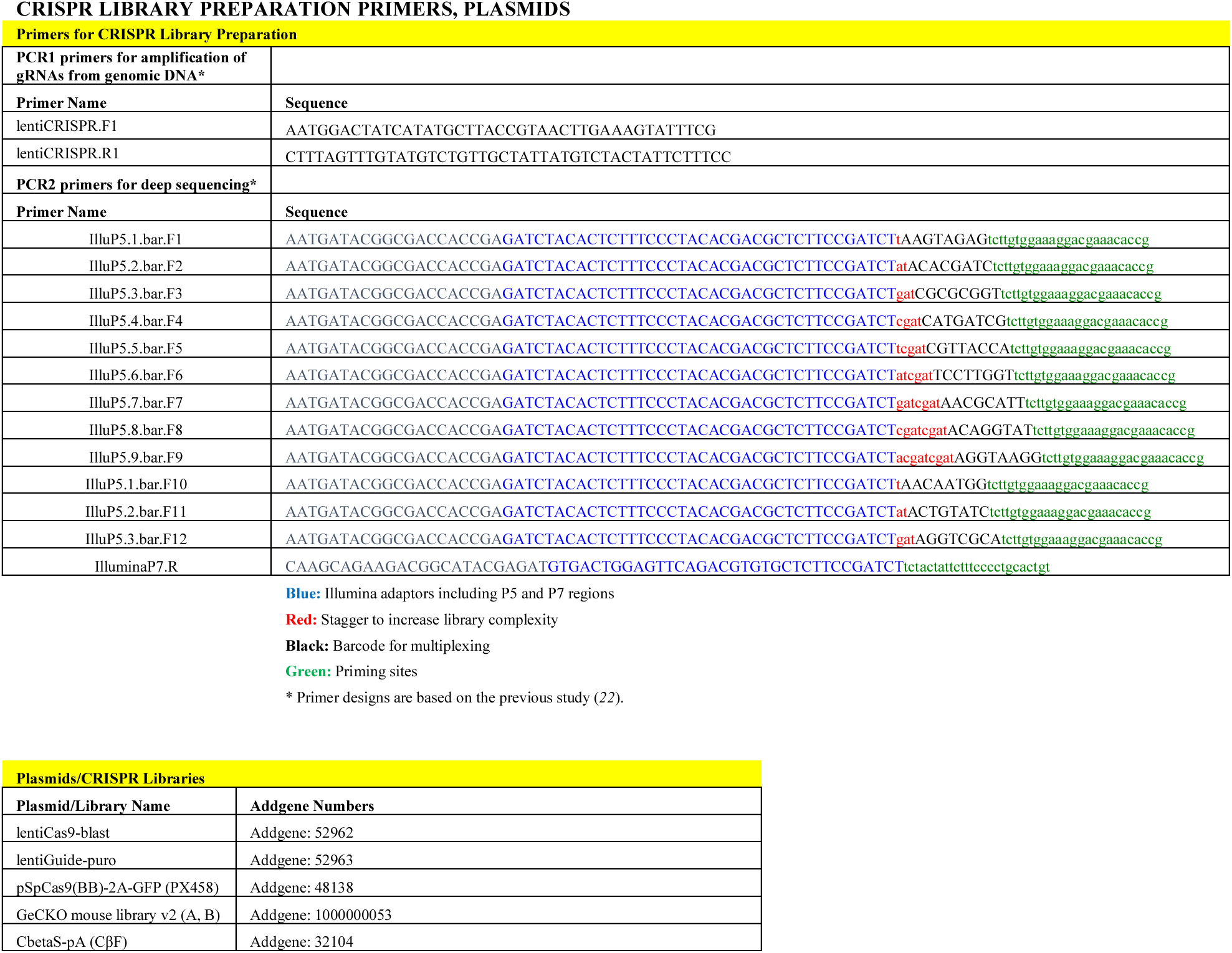
List of CRISPR library preparation primers and plasmids.

**Data S1. (separate file)**

Common candidates identified to influence CTCF boundary in independent sub-library screens

**Data S2. (separate file)**

Peptide counts in native FLAG-CTCF ChIP-MS in ESCs and MNs

**Data S3. (separate file)**

List of sgRNAs in the custom library

**Data S4. (separate file)**

List of genes identified in secondary screens in WT background

**Data S5. (separate file)**

List of genes identified in secondary screens in CTCF site deletion background

**Data S6. (separate file)**

List of genes uniquely identified in secondary screens in WT background compared to CTCF site deletion background

**Data S7. (separate file)**

RNAseq expression values in WT vs Maz KO ESCs for differentially expressed genes

**Data S8. (separate file)**

RNAseq expression values in WT vs Maz KO MNs for differentially expressed genes

**Data S9. (separate file)**

RNAseq expression values in WT vs Maz (Δ5|6) ESCs for differentially expressed genes

**Data S10. (separate file)**

RNAseq expression values in WT vs Maz (Δ5|6) MNs for differentially expressed genes

